# LEOPARD: missing view completion for multi-timepoint omics data via representation disentanglement and temporal knowledge transfer

**DOI:** 10.1101/2023.09.26.559302

**Authors:** Siyu Han, Shixiang Yu, Mengya Shi, Makoto Harada, Jianhong Ge, Jiesheng Lin, Cornelia Prehn, Agnese Petrera, Ying Li, Flora Sam, Giuseppe Matullo, Jerzy Adamski, Karsten Suhre, Christian Gieger, Stefanie M. Hauck, Christian Herder, Michael Roden, Francesco Paolo Casale, Na Cai, Annette Peters, Rui Wang-Sattler

## Abstract

Longitudinal multi-view omics data offer unique insights into the temporal dynamics of individual-level physiology, which provides opportunities to advance personalized healthcare. However, the common occurrence of incomplete views makes extrapolation tasks difficult, and there is a lack of tailored methods for this critical issue. Here, we introduce LEOPARD, an innovative approach specifically designed to complete missing views in multi-timepoint omics data. By disentangling longitudinal omics data into content and temporal representations, LEOPARD transfers the temporal knowledge to the omics-specific content, thereby completing missing views. The effectiveness of LEOPARD is validated on three benchmark datasets constructed with data from the MGH COVID study and the KORA cohort, spanning periods from 3 days to 14 years. Compared to conventional imputation methods, such as missForest, PMM, GLMM, and cGAN, LEOPARD yields the most robust results across the benchmark datasets. LEOPARD-imputed data also achieve the highest agreement with observed data in our analyses for age-associated metabolites detection, estimated glomerular filtration rate-associated proteins identification, and chronic kidney disease prediction. Our work takes the first step toward a generalized treatment of missing views in longitudinal omics data, enabling comprehensive exploration of temporal dynamics and providing valuable insights into personalized healthcare.

## Introduction

The rapid advancement of omics technologies has enabled researchers to obtain high-dimensional datasets across multiple views, enabling unprecedented explorations into the biology behind complex diseases^1^. Each view corresponds to a different type of omics data, or data acquired through a different platform, each contributing a partial or entirely independent perspective on complex biological systems^2^. While advancements in multi-omics measurements have increased throughput and enabled the acquisition of multiple views in a single assay^3^, data preprocessing, analysis, and interpretation remain significant and important challenges.

One of the most pressing challenges is the presence of missing data^4^, which at its best (when missingness occurs at random) reduces statistical power, and at its worst (when it is not random) can lead to biased discoveries. Unlike missing data points that may be scattered across the entire dataset, a missing view refers to the complete absence of all features from a certain view, as shown in Fig. 1a. In longitudinal studies that can span decades, the problem of missing views in multi-timepoint omics data becomes increasingly common due to factors such as dropout in omics measurements, experimental errors, or unavailability of specific omics profiling platforms at certain timepoints. The incompleteness of these datasets hinders multi-omics integration^5^ and investigations into predisposing factors (such as age and genetics), enabling factors (such as healthcare service and physical activity), and biomarkers for diseases.

**Fig. 1.**
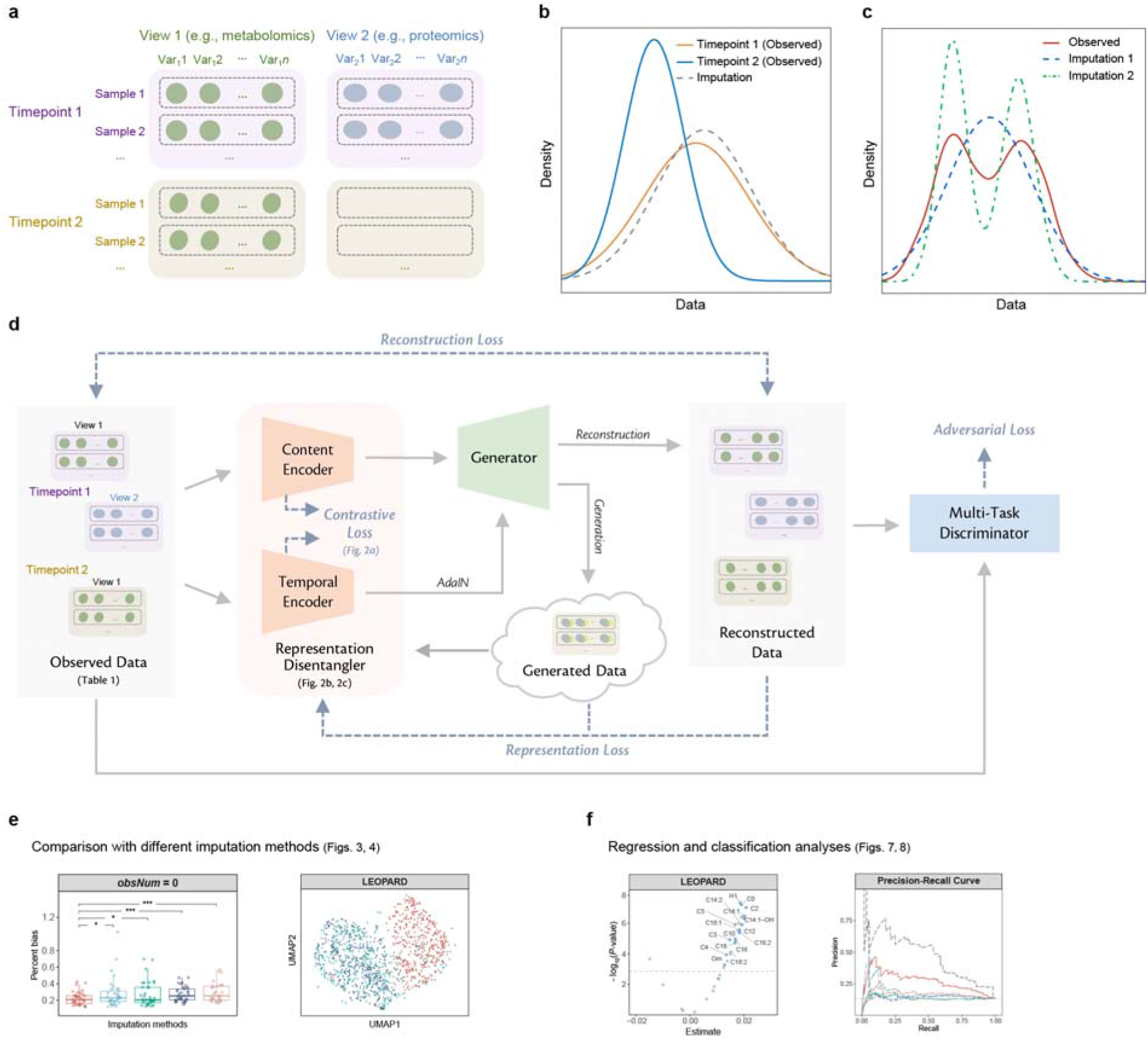
Problem description and overview of LEOPARD architecture. **a**, An example of a missing view in a longitudinal multi-omics dataset. In this instance, View 1 and View 2 correspond to metabolomics and proteomics data, respectively. Data of View 1 at Timepoint 2 are absent. **b**, An example of data density calculated from a variable in observed data (Timepoint 1 and Timepoint 2) and imputed data. The data density indicates a distribution shift across the two timepoints. Imputation methods developed for cross-sectional data cannot account for the temporal changes within the data, and their imputation models built with data from one timepoint, such as Timepoint 1, might not be appropriate for inferring data from another timepoint, such as Timepoint 2. **c**, Compared to Raw data, data of Imputation 1 may exhibit lower MSE than data of Imputation 2, but Imputation 1 potentially lose biological variations present in the data. **d**, The architecture of LEOPARD. Omics data from multiple timepoints are disentangled into omics-specific content representation and timepoint-specific temporal knowledge by the content and temporal encoders. The generator learns mappings between two views, while temporal knowledge is injected into content representation via the AdaIN operation. The multi-task discriminator encourages the distributions of reconstructed data to align more closely with the actual distribution. Contrastive loss enhances the representation learning process. Reconstruction loss measures the MSE between the input and reconstructed data. Representation loss stabilizes the training process by minimizing the MSE between the representations factorized from the reconstructed and actual data. Adversarial loss is incorporated to alleviate the element-wise averaging issue of the MSE loss. **e**, the performance of LEOPARD is evaluated with PB and UMAP embeddings. **f**, two case studies including both regression and classification analyses are performed to evaluate if biological information is preserved in the imputed data.

Missing view completion refers to the estimation of missing omics data in a multi-view context. The task of missing view completion in multi-timepoint scenario is more complex than cross-sectional missing value imputation, as it needs to harmonize data distributions across views^6^ and capture temporal patterns^7^. Generic methods, such as PMM (Predictive Mean Matching)^8^, missForest^9^, and KNNimpute^10^, learn direct mappings between views from observed data without missingness. However, this strategy is inadequate or suboptimal^11^ for longitudinal data, as it precludes investigations into temporal variation, which can be of great interest: the learned mappings may overfit the training timepoints, making them unsuitable for inferring data at other timepoints, especially when biological variations cause distribution shifts over time (Fig. 1b). To address the complexities of longitudinal data, numerous effective imputation methods have been developed based on the generalized linear mixed effect model (GLMM)^12^. Existing studies have also explored the use of spline interpolation and Gaussian processes to extrapolate or interpolate missing timepoints^13,14^. However, the typically limited number of timepoints in current human cohorts can restrict the effectiveness of these longitudinal methods. Given these challenges, there is a growing need for view completion methods that are specifically designed for multi-timepoint omics data. While metrics like mean squared error (MSE) and percent bias (PB) are commonly used to evaluate imputation results^15^, these quantitative metrics alone may not fully capture data quality in the context of omics data. As depicted in Fig. 1c, data imputed by method 1 may have a lower MSE than that imputed by method 2, but at a loss of biologically meaningful variations. Further case studies would be helpful to evaluate imputation methods.

In this paper, we introduce LEOPARD (missing view comp**l**etion for multi-tim**<u>e</u>**point **<u>o</u>**mics data via re**<u>p</u>**resentation disent**<u>a</u>**nglement and tempo**<u>r</u>**al knowle**<u>d</u>**ge transfer), a neural network-based approach that offers a novel and effective solution to this challenge (Fig.1d). LEOPARD extends representation disentanglement^16^ and style transfer^17^ techniques, which have been widely applied in various contexts such as image classification^18^, image synthesis^19^, and voice conversion^20^, to missing view completion in longitudinal omics data. LEOPARD factorizes omics data from different timepoints into omics-specific content and timepoint-specific knowledge via contrastive learning. Missing views are completed by transferring temporal knowledge to the corresponding omics-specific content.

We demonstrate the effectiveness of LEOPARD through extensive simulations using human proteomics and metabolomics data from the MGH (Massachusetts General Hospital) COVID study^21^ and the KORA (Cooperative Health Research in the Region of Augsburg) cohort^22^ (Fig. 1e). Additionally, we perform two case studies using real omics data to assess whether biological information is preserved in the imputed data, providing a comprehensive assessment of LEOPARD’s performance in both regression and classification tasks (Fig.1f).

In summary, the key contributions of this study are:

- We propose LEOPARD, a novel method tailored for missing view completion in multi-timepoint omics data that innovatively applies representation disentanglement and style transfer.
- Our study shows that generic imputation methods designed for cross-sectional data are not suitable for longitudinal data, emphasizing the need for tailored approaches. Additionally, we highlight that canonical evaluation metrics do not adequately reflect the quality of imputed biomedical data. Further investigations including regression and classification analyses can augment these metrics in assessment of imputation quality and preservation of biological variations.
- Our research reveals that omics data across timepoints can be factorized into content and temporal knowledge, providing a foundation for further explorations into biological temporal dynamics. This insight offers a novel perspective for predictive healthcare that extends beyond the problem of data imputation.

## Results

### Characterization of benchmark datasets

We evaluate LEOPARD using three real longitudinal omics datasets. These distinct datasets are designed based on data variations, time spans, and sample sizes (Table 1, Methods). The first two are mono-omics datasets, constructed with the proteomics data from MGH COVID study and the metabolomics from the KORA cohort, respectively. Views in both datasets correspond to panels or biochemical classes. Missingness in these datasets exemplifies a common issue encountered in longitudinal studies where data from certain panels or biochemical classes are incomplete in some but not all timepoints. The third dataset is a multi-omics dataset consisting of both metabolomics and proteomics data from the KORA cohort. In this dataset, views correspond to different omics. This dataset exemplifies the situation where data of a type of omics is incomplete. These three datasets comprise data of two views (*ν*1 and *ν*2 this from two timepoints (*t*1and *t*2). The samples from each dataset are split into training, validation, and test sets in a 64%, 16% and 20% ratio, respectively. We use 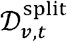 to denote data split from different views and timepoints. The test data in *ν*2 at *t*2 (i.e. 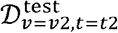) are masked for performance evaluation. To further evaluate LEOPARD’s applicability, the KORA metabolomics dataset is extended to span three timepoints, and 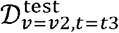 is masked for evaluation (Methods).

**Table 1.** Summary of benchmark datasets used in this study.

|  | **MGH COVID proteomics** | **KORA metabolomics** | **KORA multi-omics** |
| --- | --- | --- | --- |
| **Omics type** | proteomics | metabolomics | metabolomics; proteomics |
| **View** |  |  |  |
| $v1$ | panel: Explore 384 cardiometabolic | biochemical class<sup>*</sup>: GPL | omics: metabolomics |
| $v2$ | panel: Explore 384 inflammation | biochemical class<sup>*</sup>: AC, SL, AA, MS | omics: proteomics |
| **Variable number** | Explore 384 cardiometabolic: 322,<br>Explore 384 inflammation: 295 | GPL: 70,<br>AC: 15,<br>SL: 11,<br>AA: 9,<br>MS: 1 | metabolites: 104,<br>proteins: 66 |
| **Timepoint** |  |  |  |
| $t1$ | D0 | F4 | S4 |
| $t2$ | D3 | FF4 | F4 |
| **Time span** | 3 days | 7 years | 7 years |
| **Sample number** |  |  |  |
| Total (100%) | 218 | 2,085 | 1,062 |
| Training (64%) | 140 | 1,335 | 680 |
| Validation (16%) | 35 | 333 | 170 |
| Test (20%) | 43 | 417 | 212 |
| **Masked data for completion** | inflammation proteins from D3 | non-GPL metabolites from FF4 | proteins from F4 |

<sup>\*</sup>The targeted metabolites from KORA are classified into five analyte classes, namely glycerophospholipid (GPL), acylcarnitine (AC), sphingolipid (SL), amino acid (AA), and monosaccharide (MS). One view group comprises 70 metabolites from the glycerophospholipid class, while the other view group comprises 36 metabolites from the other classes.

### CGAN Architecture as a reference method

Existing neural network-based missing view completion methods^23–25^ have shown remarkable performance in the field of computer vision. Given their inapplicability to omics data, we designed a conditional generative adversarial network (cGAN) model specifically tailored for omics data, as a reference method. This architecture is inspired by VIGAN (View Imputation via Generative Adversarial Networks)^26^ and a method proposed by Cai *et al*.^27^, both initially designed for multi-modality image completion.

In the training phase, the generator of the cGAN is trained on observed data from the training set to capture the mappings between two views. The discriminator guides the generator to produce data with a distribution similar to that of actual data. The discriminator also has an auxiliary classifier^28^ to ensure the generated data can be paired with input data (Methods). In the inference phase, the generator utilizes the mappings it has learned from the observed data to impute the missing view in the test set. Compared to methods PMM and missForest, our cGAN model has the potential to learn more complex mappings between views. However, these three methods are not able to capture temporal changes within longitudinal data and can only learn from samples where both views are observed.

### Overview of LEOPARD architecture

Instead of relying on learning direct mappings between views, LEOPARD captures and transfers temporal knowledge to complete missing data, which also enables it to learn from samples even when only one view is available. As illustrated in Fig. 1d, the LEOPARD architecture comprises several hierarchical components. First, data of each view are transformed into vectors of equal dimensions using corresponding pre-layers. Subsequently, omics data of all views are decomposed into content and temporal representations. The content encoder captures the intrinsic content of the views, while the temporal encoder extracts knowledge specific to different timepoints. A generator then reconstructs observed views or completes missing views by transferring the temporal knowledge to the view-specific content using Adaptive Instance Normalization (AdaIN)^17^. Lastly, we use a multi-task discriminator to discriminate between real and generated data (Methods).

The LEOPARD model is trained by minimizing four types of losses: contrastive loss, representation loss, reconstruction loss, and adversarial loss. Ablation test is performed to evaluate the contribution of each loss (Methods). Minimizing normalized temperature-scaled cross-entropy (NT-Xent)-based contrastive loss^29^ optimizes the factorization of data into content and temporal representation. For both representations, minimizing the contrastive loss brings together the data pairs from the same view or timepoint and pushes apart the data pairs from different ones, so that the encoders learn similar intrinsic content (or temporal knowledge) across timepoints (or views). The representation loss, also computed on content and temporal representations, measures the MSE of the representations factorized from the actual and reconstructed data. LEOPARD minimizes this loss based on the intuition that the representations of the actual and reconstructed data should be alike. The reconstruction loss measures the MSE between imputed and observed values. Previous studies^30–32^ have demonstrated that the optimization of MSE loss often results in averaged outputs, leading to blurring effects when generating images. In our context, this might diminish biological variations present in omics data. To alleviate this issue, we use adversarial loss to encourage the predicted distribution to align more closely with the actual distribution.

LEOPARD has three unique features compared to conventional architectures for multi-view data completion. First, instead of focusing on direct mappings between views, which can only be learned from paired data where both views are present, LEOPARD formulates this imputation task in terms of representation learning and style transfer. This allows LEOPARD to utilize all available data, including observations present in only one view or timepoint. Second, it incorporates contrastive loss to disentangle the representations unique to views and timepoints, which enables the model to learn more generalized and structured representations. Our experiments show that this is of importance to improve data quality. Third, its multi-task discriminator solves multiple adversarial classification tasks simultaneously by yielding multiple binary prediction results, which has been proved to be more efficient and effective than a discriminator for a multi-class classification problem^33^.

### The representation disentanglement of LEOPARD

We use the KORA multi-omics dataset to examine if LEOPARD can effectively disentangle content and temporal representations from omics data. In this analysis, the model is trained for 600 epochs to ensure that the contrastive loss stabilizes and reaches full saturation (Fig. 2a). We use the uniform manifold approximation and projection (UMAP)^34^ for visualizing the content and temporal representations of the validation set across different views and timepoints. As the training progresses, we expect similar representations to gradually cluster together in UMAP, while dissimilar ones form distinct clusters.

**Fig. 2.**
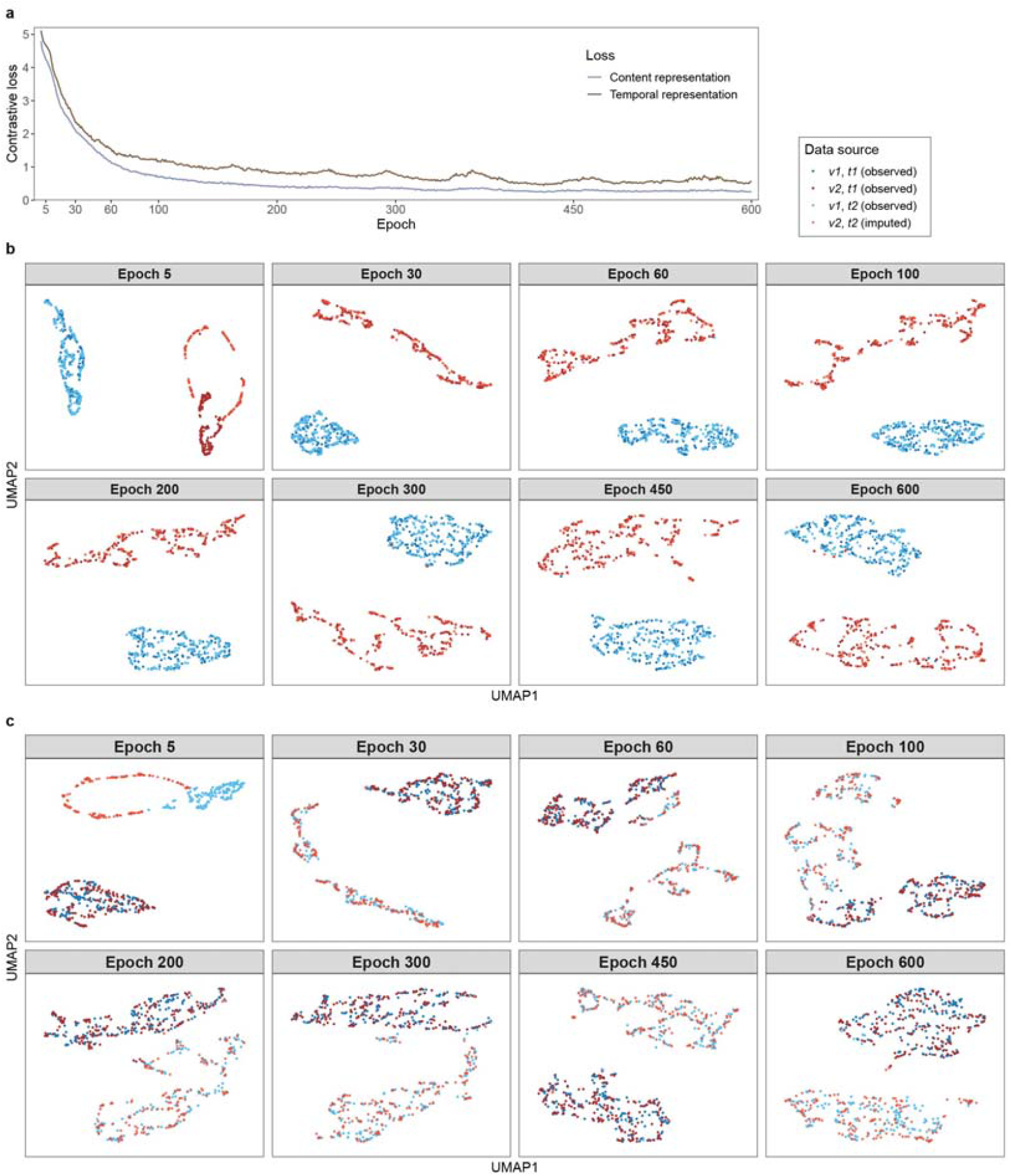
Contrastive loss and UMAP embeddings illustrate the representation disentanglement process of LEOPARD on the KORA multi-omics dataset. **a**, The NT-Xent-based contrastive loss is computed for content and temporal representations. **b-c**, UMAP embeddings of content (**b**) and temporal (**c**) representations at various training epochs are visualized for the KORA multi-omics dataset’s validation set. Representations encoded from data of *ν*1 and *ν*2 (metabolomics and proteomics, depicted by blue and red dots) at timepoint *t*1 and *t*2 (S4 and F4, depicted by dark- and light-colored dots) are plotted. The data of *ν*2 at *t*2 are imputed data produced after each training epoch, while the other data are from the observed samples in the validation set. LEOPARD’s content and temporal encoders capture signals unique to omics-specific content and temporal variations. In (**b**), as the training progresses, one cluster is formed by the data of *ν*2 at *t*1 and *t*2 (dark and light red dots), while the other cluster is formed by the data of *ν*1 at *t*1 and *t*2 (dark and light blue dots), indicating that the content encoder is able to encode timepoint-invariant content representations. Similarly, in (**c**), embeddings from the same timepoint cluster together. One cluster is formed by the data of *ν*1 and *ν*2 at t1 (dark blue and red dots), and the other is formed by the data of *ν*1 and *ν*2 at *t*2 (light blue and red dots). This demonstrates that LEOPARD can effectively factorize omics data into content and temporal representations.

As the model trains, the contrastive loss decreases (Fig. 2a), indicating that LEOPARD is increasingly able to encode the representations for different views and timepoints. The content representation embeddings (Fig. 2b) of observed *v*1(in blue) and *v*2(in red) separate rapidly during training (epoch 5), but those of the imputed *v*2for *t*2(in light red) do not mix with those of the observed *v*2 at *t*1 (in dark red). This suggests that while LEOPARD can distinguish between *v*1 and *v*2 after only a few training epochs, it is not yet capable of producing high-quality *v*2 for *t*2 that have similar content information as the observed *v*2 at *t*1. After 30 epochs of training, the content representation of *v*2 at *t*2 is better encoded, with its embeddings mixing with those of *v*2 at *t*1. Similar trends are observed in the temporal representations (Fig. 2c), where embeddings of each timepoint (*t*1 and *t*2 in dark and light colors, respectively) gradually form their corresponding clusters as training progresses. We form distinct clusters. Even after 100 epochs, some temporal representation embeddings of *t*1 notice that the temporal representations take more epochs than the content representations to are still mixed with those of *t*2. However, after around 450 epochs, LEOPARD is demonstrably able to encode temporal information that is unique to *t*1 and *t*2 (Fig. 2c).

### Benchmarking LEOPARD against conventional methods

Due to the lack of established methods specifically designed for missing view completion in multi-timepoint omics datasets, we benchmarked LEOPARD against three widely recognized generic imputation methods: missForest, PMM, and GLMM, as well as a cGAN model designed for this study. The cGAN serves as a reference model to demonstrate how existing neural network approaches, typically suited for cross-sectional data, perform in longitudinal scenarios. MissForest, as a representative non-parametric method, was chosen for its robustness and ability to handle complex, non-linear relationships among variables. PMM and GLMM, both implemented within the MICE (Multivariate Imputation by Chained Equations)^35^ framework, represent established multiple imputation methods that not only address missing values but also allow for the assessment of imputation uncertainty. GLMM, with its ability to capture temporal patterns inherent in longitudinal data, is particularly advantageous for data imputation in longitudinal scenarios.

We assess the performance of LEOPARD, cGAN, missForest, PMM, and GLMM on 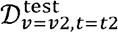 of each benchmark dataset. During training, the methods build models using data From 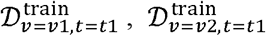, and 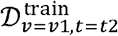, along with different numbers of training observations (*obsNum*) from 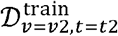. By varying *obsNum*, we assess how additional observed data from *t*2 affects their imputation performances. When *obsNum* is zero, data of *v*2 at *t*2 are assumed as completely missing, and GLMM cannot be trained due to limited longitudinal information. In this scenario, we train a linear model (LM)^36^ to complete missing view (Methods). This additionally allows us to evaluate how the performance of GLMM compares to that of the simpler LM method.

Two evaluation metrics, PB and UMAP visualization, are used for performance evaluation. PB quantifies the median absolute error ratio between the observed and imputed values for each variable, offering a variable-level assessment of imputation performance (Methods). In contrast, UMAP visualization illustrates overall similarities between observed and imputed datasets, providing a dataset-level evaluation. For the multiple imputation methods, each individual imputation is also evaluated using PB (Fig. S1) and UMAP (Fig. S2).

Apart from missing views, the presence of missing data points in observed views is also very common in omics analysis. Therefore, LEOPARD is designed to tolerate a small number of missing data points in the observed views (Methods). We further use KORA metabolomics dataset, which has the largest sample size, to evaluate the performance of different methods when observed views contain missing values. We simulate missing values by randomly masking 1%, 3%, 5%, 10%, and 20% of the data in the observed views (*maskObs*) under the assumption that data points are missing completely at random (MCAR). The experiment is repeated 10 times for each specified proportion, and the results are evaluated by PB and UMAP.

### Evaluation on the mono-omics datasets

For the MGH COVID proteomics dataset, missForest overall exhibits the lowest PB, whereas LEOPARD performs slightly worse than missForest and its neural network-based counterpart, cGAN (Fig. 3, upper row). When compared to LM, GLMM does not show improved performance. As *obsNum* increases, the PB values of all methods tend to decrease, and the performance gap between LEOPARD and missForest diminishes. Specifically, when *obsNum* is 100, the UMAP representation (Fig. 4, upper row) reveals that the clusters of the imputed data generated by all five methods (green dots) closely approximated the actual data (blue dots), indicating high similarity between the imputed and original datasets.

**Fig. 3.**
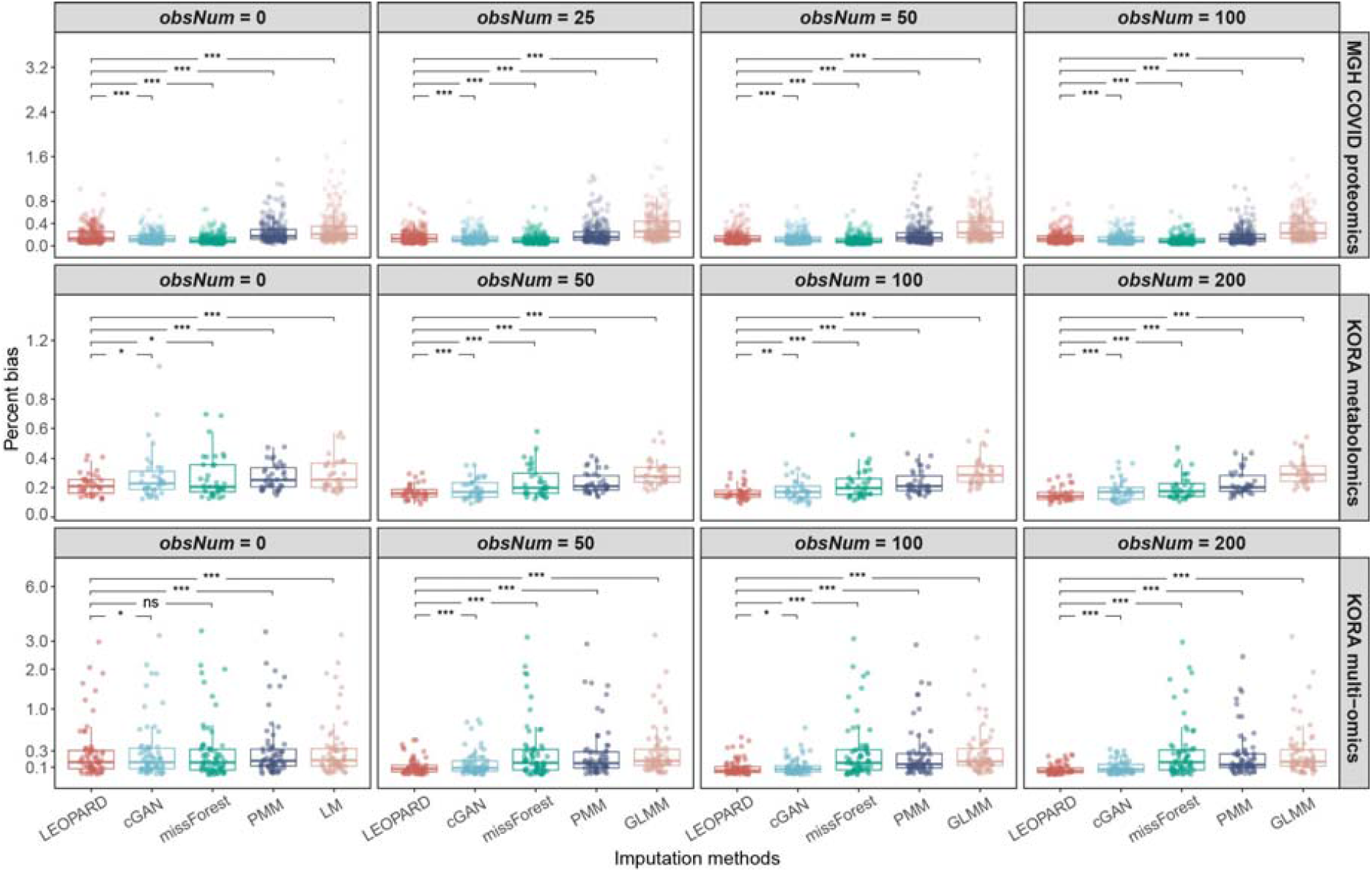
PB of imputed results for the test sets of three benchmark datasets. PB evaluated on 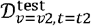 of the MGH COVID proteomics dataset (upper row), KORA metabolomics dataset (middle row), and KORA multi-omics dataset (lower row). Each dot represents a PB value for a variable. Please note that LM is used for imputation instead of GLMM when *obsNum* = 0. Significance level: not significant (ns), *P* < 0.05 (*), *P* < 0.01 (**), and *P* < 0.001 (***).

**Fig. 4.**
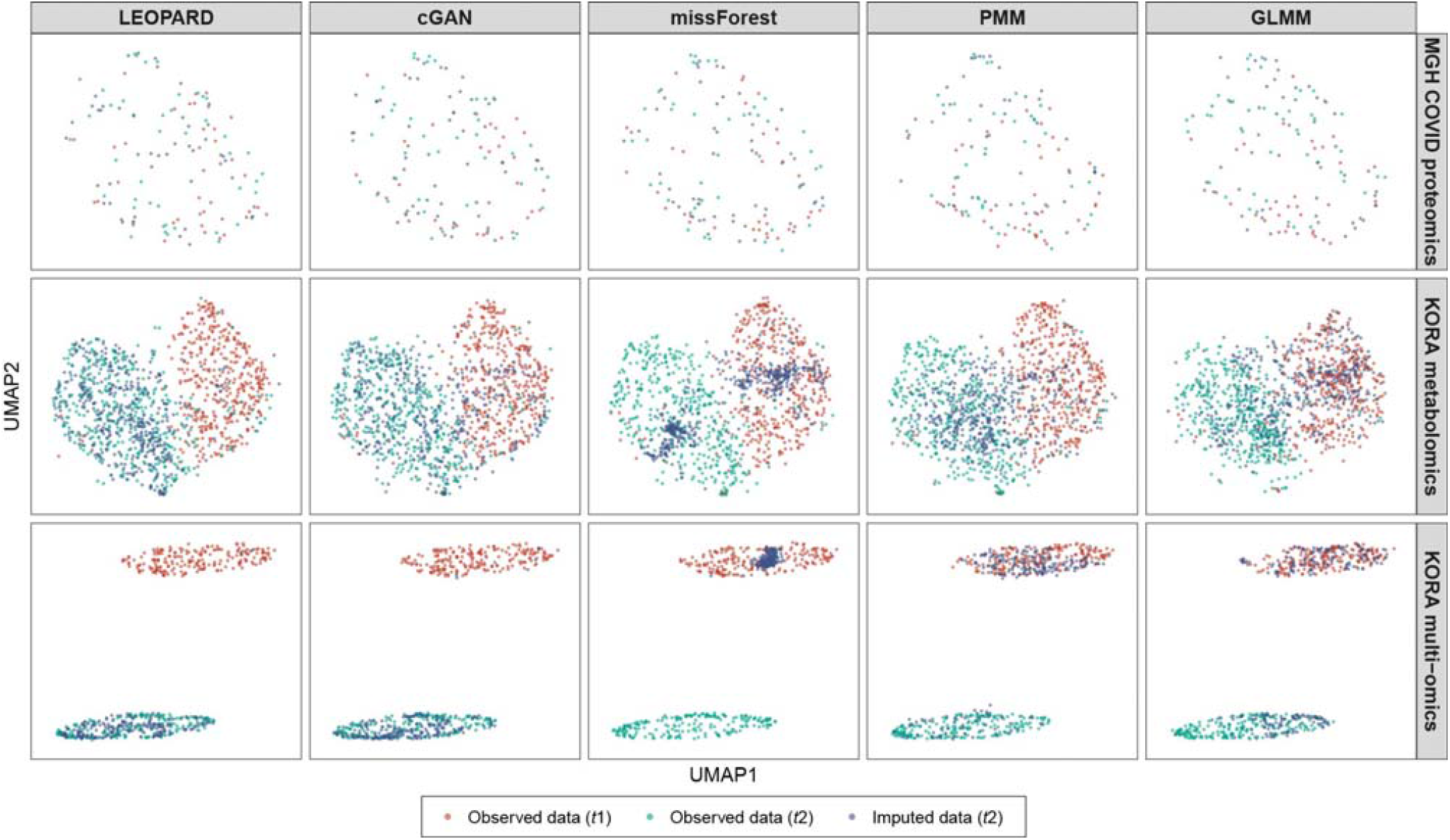
UMAP representations of the imputed values and corresponding observed data from the benchmark datasets. UMAP models are initially fitted with the training data from the MGH COVID proteomics dataset (upper row, *t*1: D0, *t*2: D3), KORA metabolomics dataset (middle row, *t*1: F4, *t*2: FF4), and KORA multi-omics dataset (lower row, *t*1: S4, *t*2: F4). Subsequently, the trained models are applied to the corresponding observed data (represented by red and blue dots for *t*1 and *t*2) and the data imputed by different methods (represented by green dots) under the setting of *obsNum* = 100 for the MGH COVID dataset and *obsNum* = 200 for the two KORA-derived datasets. The distributions of red and blue dots illustrate the variation across the two timepoints, while the similarity between the distributions of blue and green dots indicates the quality of the imputed data. A high degree of similarity suggests a strong resemblance between the imputed and observed data.

Interestingly, we observe that missForest, despite yielding the best performance for the MGH COVID dataset, produces the most unstable result for the KORA metabolomics dataset, showing the largest interquartile range (IQR) of 0.186 when *obsNum* is 0 (Fig. 3, middle row). In comparison, LEOPARD achieves the smallest IQR of 0.094 under the same condition, while cGAN, PMM, and LM obtain IQR values of 0.125, 0.132, and 0.166, respectively. As *obsNum* increases to 200, LEOPARD, cGAN, missForest, and PMM lower their median PB values to 0.142, 0.172, 0.177, and 0.204, respectively. However, GLMM obtains a median PB of 0.291 and does not outperform LM. From the UMAP plots generated from the data imputed under *obsNum* = 200, we notice a large amount of the embeddings from 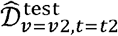 generated by cGAN, missForest, and GLMM are mixed with those of 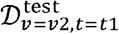, instead of 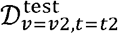, implying that they overfit to *t*1 and do not generalize well to the second timepoint (Fig. 4, middle row). Moreover, the UMAP embeddings of 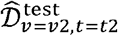 from missForest and PMM only partly overlap with those of 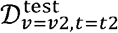, suggesting that some variations in the observed data have not been captured. In contrast, the embeddings of the data imputed by LEOPARD widely spread within the embedding space of the observed data, which demonstrates that LEOPARD has effectively learned and approximated the observed data’s distribution.

### Evaluation on the KORA multi-omics dataset

In contrast to mono-omics datasets, where both views are from the same omics type, multi-omics datasets require imputation methods to capture more intricate relationships between omics data to ensure accurate results. In our evaluation of the KORA multi-omics data, all methods show some extremely high PB values when *obsNum* is 0 (Fig. 3, lower row). As *obsNum* increases to 200, LEOPARD greatly reduces its median PB from 0.152 to 0.061, outperforming its closest competitor, cGAN, which reduces its median PB from 0.158 to 0.076. In contrast, the performances of missForest (from 0.156 to 0.159) and PMM (from 0.177 to 0.131) do not show similar improvements. GLMM reduces its median PB from 0.176 to 0.163 as *obsNum* increases from 50 to 200. The UMAP visualization further reveals a limited ability of missForest, PMM, and GLMM to capture signals from the *t*2 timepoint, as their embeddings of 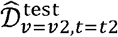 cluster with 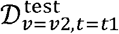, not 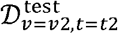 (Fig. 4, lower row). LEOPARD’s performance is further validated by a high similarity between the distributions of the imputed and observed data embeddings in the UMAP space.

### Analysis on imputed data with extremely high PB values

We then investigate the extremely high PB values (> 0.8) observed in the KORA multi-omics dataset. Under *obsNum* = 0, we notice that proteins with low abundance (< 4.0) tend to exhibit extremely high PB in the imputed values (Fig. 5). For instance, SCF (stem cell factor), with a median abundance of 9.950, has a PB of 0.090 calculated from the LEOPARD-imputed data. In contrast, NT3 (Neurotrophin-3), with a much lower median abundance of 0.982, shows a PB of 1.187 calculated from the same imputed data. Increasing *obsNum* can substantially lower these extremely high PB values for LEOPARD and cGAN, but makes no similar contributions for missForest, PMM, and GLMM.

**Fig. 5.**
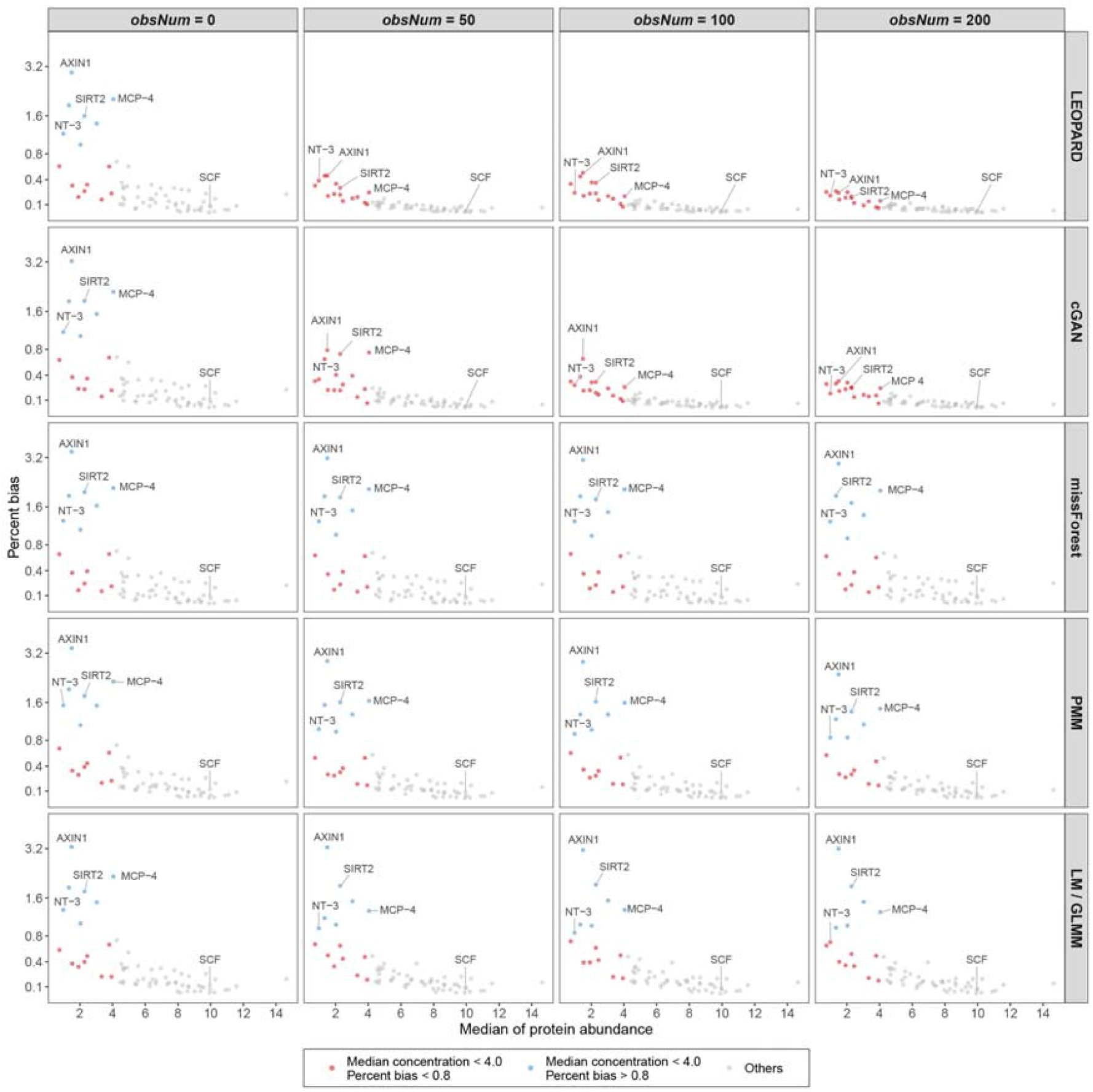
Proteins with low abundance tend to exhibit high PB in the imputed values. The proteins with low abundance (median concentration < 4.0) tend to exhibit extremely high PB (> 0.8) in the imputed values obtained under *obsNum* = 0. The extremely high PB values of LEOPARD can be lowered by increasing *obsNum*. Please note that LM is used for imputation instead of GLMM when *obsNum* = 0.

### Evaluation on observed views with missing data points

We further evaluate how different methods perform when observed views contain missing values. The PB values are averaged across 10 repetitions. The UMAP plots visualize the repetition that exhibits the lowest median of PB.

Our findings indicate that all the methods experienced an increase in PB when the observed views contain missing values (Extended Fig. 1). However, LEOPARD and missForest are robust to the missing data points in terms of PB. In contrast, cGAN and GLMM exhibit high sensitivity to those missing values. Method cGAN does not show similar improvement with the increase of *obsNum* as it performs in Fig. 3 (middle row) and is gradually surpassed by missForest. GLMM overall exhibits higher PB than the other methods.

The UMAP plots (Extended Fig. 2) further demonstrate that LEOPARD’s performance remains comparable to scenarios with no missing data in the observed views (Fig. 4, middle row), unlike the other methods which display overfitting or a great loss of data variation. Although LEOPARD outperforms other methods, we observed a change in the distribution of the imputed data (blue dots): as *maskObs* increases, these blue dots begin to shrink toward their center and become more concentrated. This leads to a reduced coverage of the outer areas of the ground truth embeddings (green dots) and suggests that the imputed data might not capture the full variability of the data when the proportion of missing data is high.

### Case studies

We perform several case studies, covering both regression and classification tasks, to investigate whether biological signals are preserved in the imputed data obtained at *obsNum* = 0.

### Regression analysis

The regression models are fitted using the observed data and different imputed data corresponding to 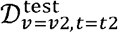. The performance of LEOPARD, cGAN, and missForest are evaluated on their imputed data directly, while two multiple imputation methods, PMM and LM, are evaluated by pooling their multiple estimates using Rubin’s rules^36^ (Methods).

We use the KORA metabolomics dataset to identify metabolites associated with age, controlling for sex. The models are fitted separately to each of the 36 metabolites (*N* = 417). Of the 18 metabolites significantly associated with age after a Bonferroni correction for multiple testing (*P* < 0.05/36) in the observed data, 17 are also significant in the data imputed by LEOPARD (see Fig. 6a). Among these 17 metabolites, several, including C14:1 (Tetradecenoylcarnitine), C18 (Octadecanoylcarnitine), C18:1 (Octadecenoylcarnitine), and Orn (Ornithine), have been validated by previous research^37–40^ showing that they might be particularly relevant in aging and age-related metabolic conditions. In contrast, only one metabolite is significantly associated with age in the data imputed by missForest. No metabolite is identified as significant in the data imputed by cGAN, PMM, and LM. The results on each imputation of PMM and LM are shown in Fig. S3.

**Fig. 6.**
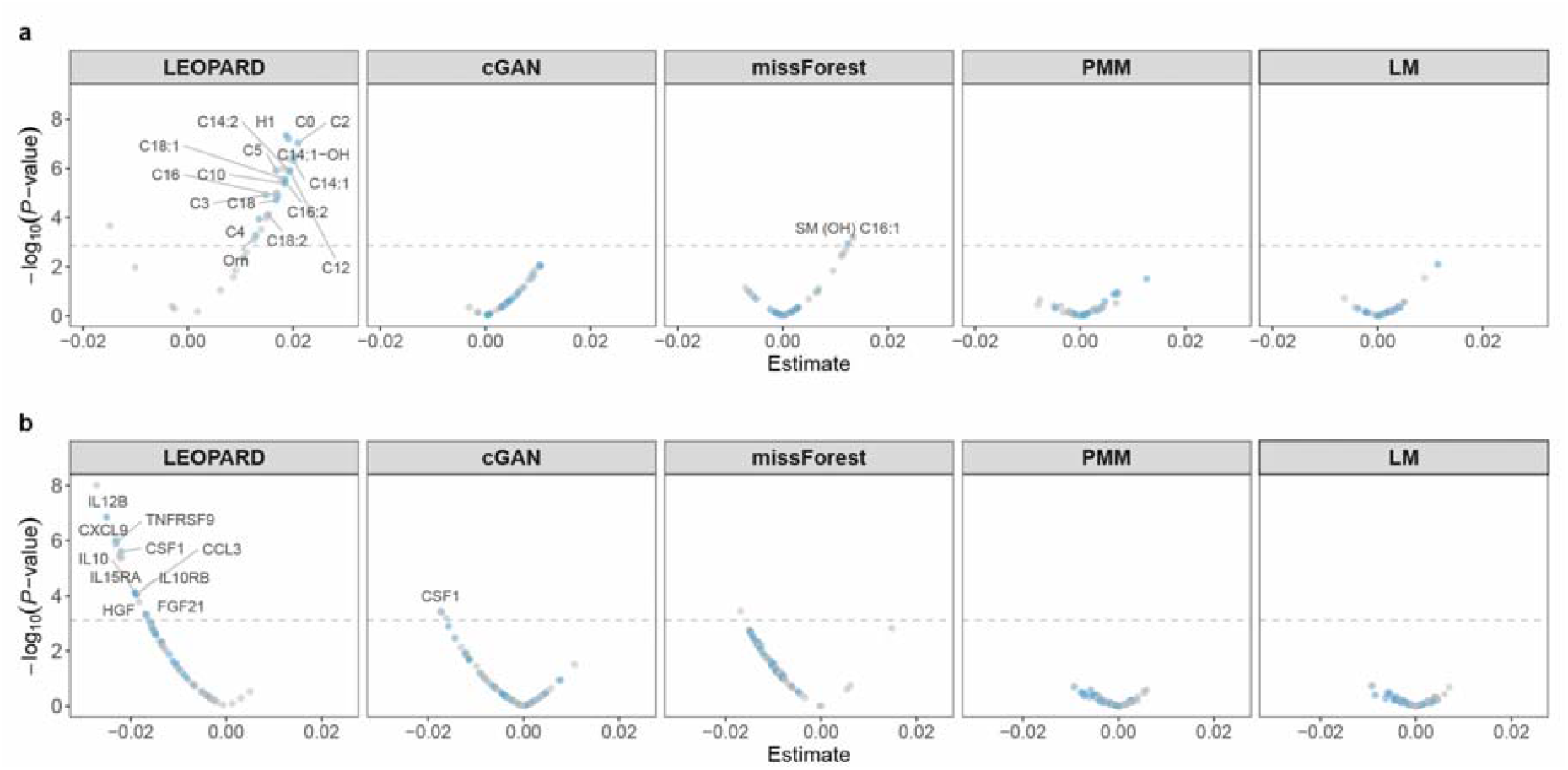
Regression analyses with the data imputed by different methods. **a**, Volcano plots display age-associated metabolites detected in the 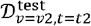 and 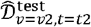 (*obsNum* = 0) of the KORA metabolomics dataset (*N* = 417). 18 significant metabolites (P < 0.05/36) identified in the observed data are shown in blue. Replicated metabolites from the data imputed by different methods are marked with labels. **b**, Volcano plots display eGFR-associated proteins detected in the 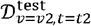 and 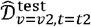 (*obsNum* = 0) of the KORA multi-omics dataset (*N* = 212). 28 significant metabolites (P < 0.05/66) identified in the observed data are shown in blue. Replicated metabolites from the data imputed by different methods are marked with labels.

We then use the KORA multi-omics dataset to identify proteins associated with the estimated glomerular filtration rate (eGFR), controlling for age and sex. Each model is individually fitted to one of the 66 proteins (*N* = 212). In the observed data, 28 proteins are significantly associated with eGFR after a Bonferroni correction (*P* < 0.05/66). Of these 28 proteins, 10 proteins remain significant in the data imputed by LEOPARD (see Fig. 6b), while one is significant in the data from cGAN, and none is identified as significant in the data from missForest, PMM, and LM. Among the 10 proteins detected from the LEOPARD-imputed data, eight (TNFRSF9, IL10RB, CSF1, FGF21, HGF, IL10, CXCL9, IL12B) have been validated by prior research^41^. The results on each imputation of PMM and LM are shown in Fig. S4.

### Classification analysis

We use balanced random forest (BRF)^42^ classifiers to predict chronic kidney disease (CKD) using the KORA metabolomics and multi-omics datasets. The classifiers are individually fitted using the observed and different imputed data of 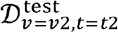, corresponding to 36 metabolites from the KORA metabolomics dataset (*N* = 416, one sample removed due to a missing CKD label) and 66 proteins from the KORA multi-omics dataset (*N* = 212). CKD cases are defined as having an eGFR < 60 mL/min/1.73m^2 43^. In the two datasets, 56 and 36 individuals are identified as CKD cases, respectively. We train the classifiers using identical hyperparameters and use leave-one-out-cross-validation (LOOCV) to evaluate their performance. The models for LEOPARD, cGAN, and missForest are trained using their respective imputed data, while the models for PMM and LM are trained on the average estimates across their multiple imputations (Methods).

For the KORA metabolomics dataset, the observed data obtain an F1 Score of 0.439, and the data imputed by LEOPARD achieves the closest performance with an F1 Score of 0.358 (Fig. 7 a, Extended Table 1). LEOPARD also outperforms its competitors in terms of accuracy, sensitivity, precision, AUROC (area under the receiver operating characteristic curve) and AUPRC (area under the precision-recall curve). The proteins from the KORA multi-omics dataset perform better than the metabolites from the metabolomics dataset for this task. The F1 Score increases to 0.544 for the observed data of the KORA multi-omics dataset. LEOPARD outperforms its competitors with an F1 Score of 0.403, an AUROC of 0.725, and an AUPRC of 0.435 (Fig. 7b, Extended Table 2). The prediction results on each individual imputation of PMM and LMM are displayed in Figs. S5 and S6.

**Fig. 7.**
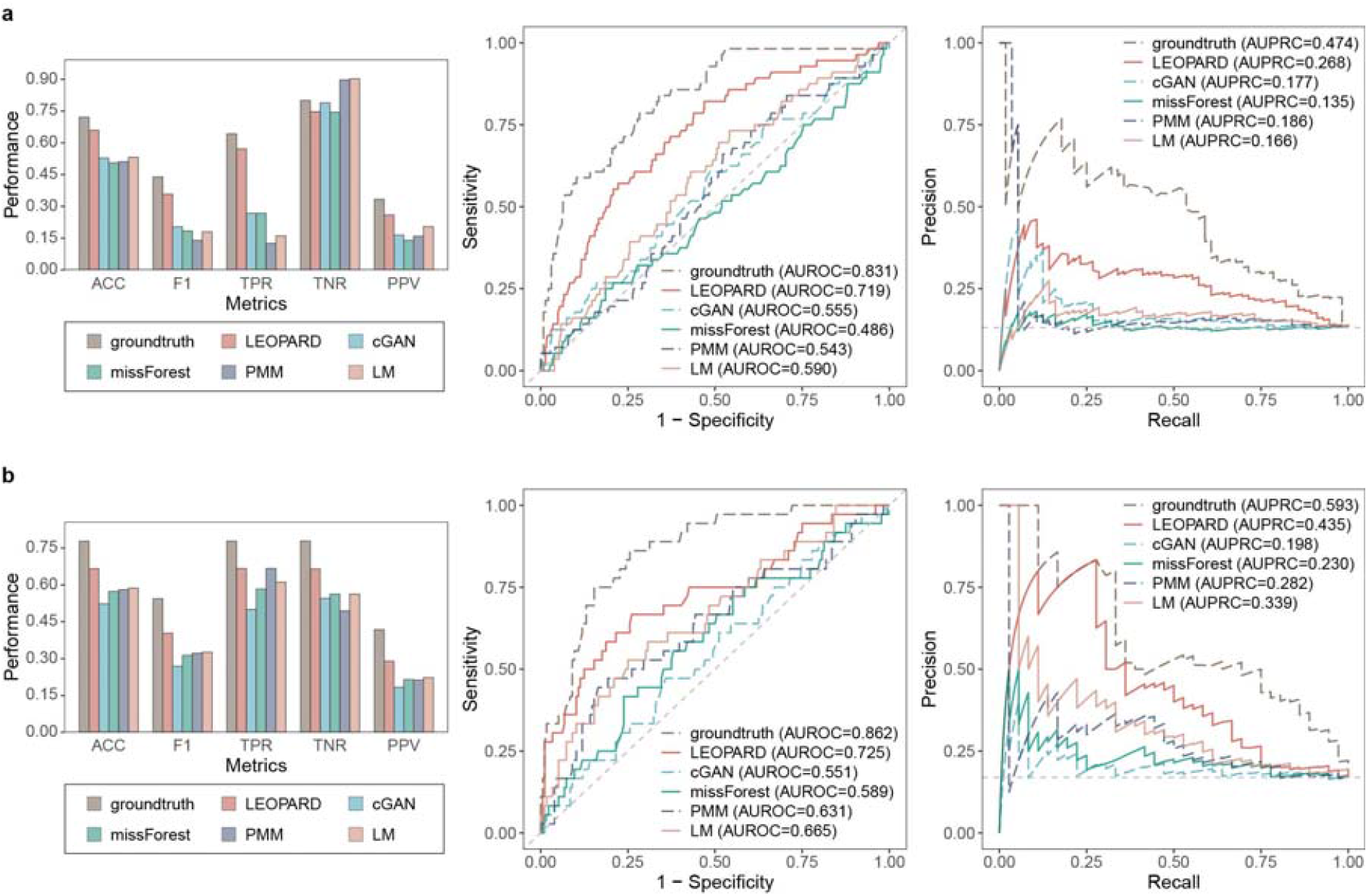
Classification analyses with the data imputed by different methods. CKD classification evaluated using 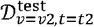 and 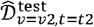 (*obsNum* = 0) from (**a**) the KORA metabolomics dataset (*N* = 416, *N*_*positive*_ = 56, *N*_*negative*_ = 360) and (**b**) the KORA multi-omics dataset (*N* = 212, *N*_*positive*_ = 36, *N*_*negative*_ = 176). Models are trained using the BRF algorithm with identical hyperparameters and evaluated using LOOCV. Evaluation metrics in the bar plot include true positive rate (TPR, also known as sensitivity), true negative rate (TNR, also known as specificity), positive predictive value (PPV, also known as precision), accuracy (ACC), and F1 score. The dashed lines in the ROC and PR curves represent the performance of a hypothetical model with no predictive capability.

### Evaluation of the applicability of LEOPARD

We next explore how many training samples are required for LEOPAR to have robust view completion and assess the performance of LEOPARD in completing different missing views at different timepoints. These analyses reveal LEOPARD’s utility and adaptability in different analytical scenarios.

#### Minimum training samples for robust view completion

The evaluation is performed on 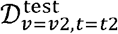 of our three benchmark datasets by varying the number of training samples from 20 to 160 and *obsNum* from 0 to 50. Each condition is tested 10 times with different samples randomly selected from the training sets. The performance is evaluated by PB averaged across these repetitions. Fig. 8 simplifies the boxplot and shows the median and the IQR of the averaged PB values calculated for the variables in the imputed data.

**Fig. 8.**
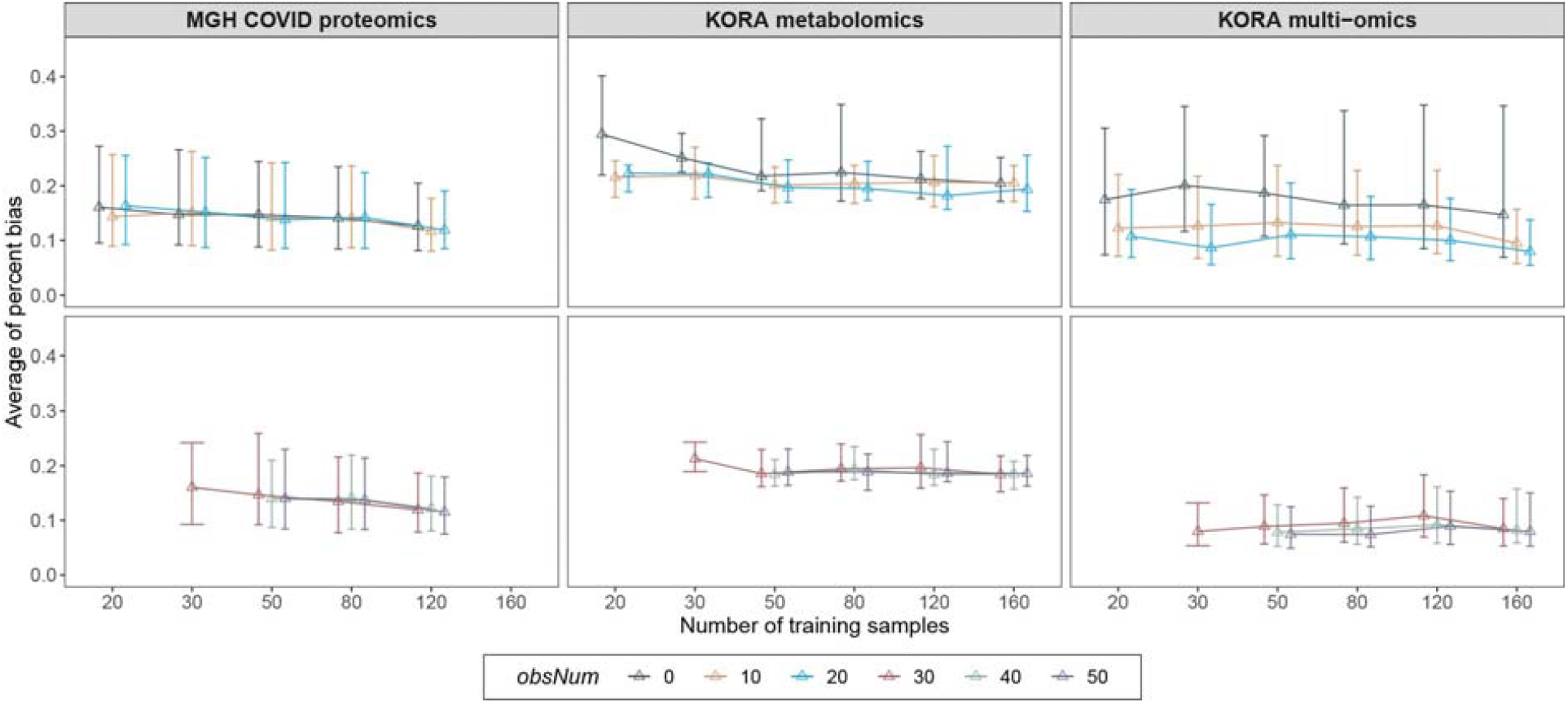
Evaluation of minimum number of training samples required for LEOPARD. For each benchmark dataset, the average PB is evaluated on 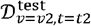 across 10 repeated completions for each combination of training sample sizes and *obsNum*. The bar indicates the median and the IQR of the average PB values for different variables. In each repetition, the samples are selected randomly. Please note that the maximum *obsNum* cannot exceed the number of training samples, and the full training set of the MGH COVID proteomics dataset contains only 140 samples.

Across all datasets, average PB generally decreases with more training samples, indicating an improvement in view completion. Consistent with our previous evaluation, PB values decrease as *obsNum* increases. Additionally, we notice that the average PB steadily decreases for the MGH COVID proteomics dataset, which exhibits the smallest variation between the two timepoints in our UMAP plots (Fig. 4). In contrast, the average PB for the other two datasets shows some fluctuations, particularly for the KORA multi-omics dataset, which shows the most obvious variation between two timepoints. When *obsNum* = 0, the MGH COVID proteomics and the KORA metabolomics datasets require about 120 training samples to obtain stable results; the KORA multi-omics dataset, however, exhibits a wide range of PB under this condition. When we increase *obsNum* to 20, the performance stabilizes with approximately 60 to 80 samples used for training LEOPARD. Based on our evaluation, at least 80 training samples may be required for robust view completion.

#### Arbitrary temporal knowledge transfer

We finally use LEOPARD to separately complete 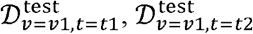, and 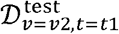 in the KORA metabolomics dataset (F4 as *t*1, FF4 as *t*2, see Table 1). Completing 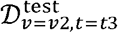 using the data from S4 (*t*1), and FF4 (*t*3) is also evaluated as a complementary analysis to our previous evaluation (Methods).

Our results reveal that some metabolites exhibit high PB values at *obsNum* = 0 (Extended Fig. 3) due to their low concentrations. While different completions show variability in their performances, PB generally decreases as *obsNum* increases. This evaluation demonstrates that LEOPARD can transfer extracted temporal knowledge to different content representations in a flexible and generalized way. However, additional observations from the incomplete view may be necessary to ensure robust results, particularly for metabolites with low concentrations.

## Discussion

In this study, we introduce LEOPARD, a novel architecture for missing view completion designed for multi-timepoint omics data. The performance of LEOPARD is comprehensively assessed through simulations and case studies using three real human omics datasets. Additional interesting findings emerged from our evaluation.

As illustrated in the UMAP plots, the MGH COVID proteomics data (Fig. 4a) from D0 (*t*1, in red) and D3 (*t*2, in blue) show relatively low variation, while the KORA metabolomics data (Fig. 4b) from F4 (*t*1, in red) and FF4 (*t*2, in blue) exhibit more substantial variations, potentially due to biological variations spanning seven years and technical variation from different analytical kits in the KORA data. In the MGH COVID dataset, we observed that LEOPARD performs slightly worse than its competitors. This can be attributed to the inherent differences in the representation learning process of LEOPARD and its competitors. By directly learning mappings between views, the methods developed for cross-sectional data can exploit the input data to learn detailed, sample-specific patterns, while the representation learning in LEOPARD primarily focuses on more compact and generalized structures related to views and timepoints, potentially neglecting detailed information specific to individual samples. The MGH COVID dataset, having a high similarity between the data from D0 and D3, allows the cross-sectional imputation methods to effectively apply the mappings learned from one timepoint to another. As the data variation between two timepoints increases, LEOPARD’s advantages become increasingly evident, while its competitors tend to overfit the training data and fail to generalize well to the new timepoint (Fig. 4, middle and lower rows).

For the KORA multi-omics dataset, we observed that the extremely high PB values are associated with low analyte abundances. For the same absolute error in imputed values, a variable with a low analyte abundance will have a higher absolute error ratio, leading to a higher PB than those variables with high abundances. Additionally, protein levels quantified using the Olink platform are represented as relative quantities. Data from different measurements also contain technical variations that arise from experimental factors and normalization methods used for relative quantification. When *obsNum* is 0, LEOPARD is trained without any data from *v*1 at *t*2, and thus cannot account for the technical variations exclusive to that part. By incorporating a few observed samples from the second timepoint into the training process, the model can better capture the data distribution and technical variation of the missing part, which contributes to a substantial reduction in high PB values.

As data imputation inevitably incurs a loss of information, we conducted case studies to assess the preservation of biological information in the imputed data. Despite all five imputation methods producing similar PB when *obsNum* is 0 (Fig. 3 lower row), the case studies showed that the data imputed by LEOPARD provided performance closest to the observed data, while the data imputed by cGAN, missForest, PMM, and LM showed a substantial loss of biological information (Fig. 5). This outcome highlights the importance of case studies for a reliable evaluation of imputed data.

Arbitrary style transfer, a concept from the computer vision field underpinning LEOPARD, allows the style of one image to be transferred to the content of another. This study demonstrates that LEOPARD inherits this capability and has the potential for arbitrary temporal knowledge transfer. Our experiments also demonstrate that LEOPARD can yield robust results with approximately 80 samples. Moreover, LEOPARD not only completes missing views for downstream analyses, but also facilitates the exploration of temporal dynamics. By analyzing the extracted temporal embeddings, LEOPARD could enable the inference of the temporal ordering of omics changes, which would be particularly valuable when there is a discrepancy between biological and chronological order. As the number of data timepoints increases, LEOPARD is expected to offer new opportunities in predictive healthcare with multi-timepoint omics data.

While LEOPARD demonstrates superior performance over existing generic imputation methods on missing view completion, it is important to consider the limitations and caveats of this study. To align with real-world settings, we defined QC criteria based on existing studies when constructing our benchmark datasets, and consequently, only the most detectable proteins and metabolites were selected. This could inflate the metrics of both LEOPARD and other methods reported in this study. The performance on these selected variables may not accurately reflect that of the overall proteins and metabolites, especially those showing more variability in their abundance. LEOPARD typically requires observed views to be complete so that temporal and content representations can be extracted. Considering the common occurrence of missing values in real-world omics data, LEOPARD is designed to tolerate a small proportion while maintaining optimal robustness. However, we observed that LEOPARD struggles to capture the full diversity present in the ground truth as *maskObs* increases to 20% (Extended Fig. 2). It is preferable for the input data for LEOPARD to contain less than 10% missing data points. Higher proportions of missing values are ideally addressed by generic imputation methods before processing with LEOPARD. Additionally, we assumed that the missing data were MCAR. Additional bias could be introduced if data points are missing at random (MAR) or missing not at random (MNAR) in real-world scenarios. Finally, our experiments were restricted by data availability to three timepoints, but in principle LEOPARD can accommodate additional timepoints and is well-suited for analyses involving multiple omics per timepoint.

With advancements in omics measurement technology and the growing availability of longitudinal data, missing view in multi-view, multi-timepoint data is becoming a prominent issue. Our study demonstrates that established generic methods, originally developed for missing data points or cross-sectional data, do not produce robust results in this new context. This highlights the necessity for specialized methods, and our method LEOPARD represents an early attempt to address this issue. We anticipate further developments in imputation methods that exhibit high generalization ability, robustness to low analyte abundance, and preservation of biological variations.

## Methods

### Terminology

The following outlines the terminology used throughout this study.

- Multi-view: different representations extracted from different data sources or several modalities collected in one cohort^2^. It is common for samples to be examined from multiple perspectives using different platforms or technologies, yielding different views of the data.
- Mapping: the transformation that a model learns to apply from an input space to an output space.
- Embeddings: low-dimensional representations of high-dimensional data.
- Arbitrary style transfer: a technique in computer vision field that allows the visual style from any source image to a target image, irrespective of their content differences. LEOPARD adapts this technique and has the potential to perform arbitrary temporal knowledge transfer.
- Representation disentanglement: the process of decomposing complex data into distinct underlying factors of variation, where each factor is represented independently of the others.

### Problem Formulation

In our missing view completion problem, we are given a generalized dataset 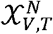 that includes omics data from *N* individuals across multiple views (*V*) and timepoints (*T*). For simplification, we consider data from two views {*v*1,*v*2}and two timepoints{ *t*1, *t*2}. Data in 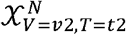 are set to be incomplete. The dataset is split into training, validation, and test sets, denoted as 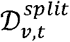. The task is to complete 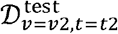. To achieve this, we attempt to develop a model that captures the mappings between *v*1 and *v*2, while simultaneously extracting temporal knowledge from *t*1 and *t*2 using all observed data in the training set.

### Data preprocessing

#### MGH COVID dataset

The MGH COVID study includes plasma proteins measured for patients at three timepoints: on day 0 (D0) for all patients on the day they were hospitalized for COVID, and on days 3 (D3) and 7 (D7) for patients still hospitalized then. We utilize proteomics data from D0 (*t*1) and D3 (*t*2), which have the largest sample sizes (*N* = 218, one duplicated sample removed), to construct the first mono-omics dataset.

The proteomics data from the MGH COVID study were obtained using plasma samples and the Olink^®^ Explore 1536 platform (Olink Proteomics, Watertown, MA, USA), which consists of 1,472 proteins across four Olink^®^ Explore 384 panels: inflammation, oncology, cardiometabolic, and neurology^21^. The platform enables relative quantification of analyte concentrations in the form of log_2_-scaled normalized protein expression (NPX) values, where higher values correspond to higher protein levels. We selected proteins listed in the Cardiometabolic and Inflammation panels to construct the two views for the MGH COVID proteomics dataset, and the inflammation view was assumed to be incomplete. The proteins are processed based on the following quality control (QC) criteria: (1) no missing values; (2) at least 50% of measured sample values are equal to or above the limits of detection (LOD). After quality inspection (Table S1), a total of 322 and 295 proteins from the Cardiometabolic (*v*1) and Inflammation (*v*2) panels were selected.

#### KORA datasets

Data from the KORA study are extracted from the baseline survey (S4, examined between 1999 and 2001), the first follow-up study (F4, 2006∼2008), and the second follow-up study (FF4, 2013∼2014)^44,45^. We use metabolomics data (*N* = 2,085) from F4 (*t*1) and FF4 (*t*2) from the KORA cohort to construct the second mono-omics dataset. We additionally use metabolomics and proteomics data (*N* = 1,062) from the KORA S4 (*t*1) and F4 (*t*2) to construct the multi-omics dataset.

The metabolite profiling of the KORA S4 (March - April 2011), F4 (August 2008 - March 2009), and FF4 (February - October 2019) serum samples spans over a decade during which analytical procedures have been upgraded several times. The targeted metabolomics data of F4 were measured with the analytical kit AbsoluteIDQ^®^ p150, while S4 and FF4 data were quantified using the kit AbsoluteIDQ^®^ p180 (Biocrates Life Sciences AG, Innsbruck, Austria). To assess the technical variation introduced by different kits, samples from 288 individuals from the F4 were remeasured (September-October 2019) using the p180 kit. Three manufacturer-provided QC samples were added to each plate to quantity the plate effect. Additionally, five external QC samples were added to each plate when measuring using the p180 kit. The p150 and p180 kits allow simultaneous quantification of 163 and 188 metabolites, respectively^46^. Only metabolites meeting the following QC criteria^47,48^ were selected: (1) overlap between p150 and p180; (2) no missing values; (3) at least 50% of measured sample values are equal to or above the LOD of corresponding plates; (4) median relative standard deviation (RSD) of QC samples < 25%; (5) Spearman correlation coefficients between the KORA F4 (remeasured, p180) and F4 (original, p150) > 0.5. After QC procedures, the metabolites values were further normalized using TIGER (Technical variation elImination with ensemble learninG architEctuRe)^46^ with its default setting to remove the plate effects. For the multi-omics dataset, TIGER was also used to remove the technical variation introduced by different kits following our previous protocol^49^.

The proteomics data from the KORA cohort are available at two timepoints, S4 and F4, and are measured using plasma (S4, February 2020) and serum (F4, December 20–6 - January 2017) samples with the Olink^®^ Target 96 Inflammation panel (Olink Proteomics, Uppsala, Sweden)^41^. The panel includes 92 proteins, and only proteins pass the following QC criteria were selected: (1) no missing values; (2) at least 75% of measured sample values are equal to or above the LOD. TIGER was then used to remove the technical variation introduced by different kits following our previous protocol^46,49^.

For the KORA metabolomics dataset, 106 targeted metabolites satisfy all criteria (Table S2) and are categorized into five analyte classes: acylcarnitine (AC), amino acid (AA), glycerophospholipid (GPL), sphingolipid (SL), and monosaccharide (MS). Two view groups are constructed by 70 metabolites from GPL (*v*1) and 36 metabolites from the other four classes (*v*2). For the KORA multi-omics dataset, 104 metabolites (*v*1) and 66 proteins (*v*2) satisfy all QC criteria (Table S2 and S3) and are selected to construct two views.

To evaluate LEOPARD’s capability for arbitrary temporal knowledge transfer, we further expanded the KORA metabolomics dataset to include data from the baseline study (S4, as *t*1) and the second follow-up study (FF4, as t3), spanning approximately 14 years. We divided the metabolites data into two views using the same strategy as we used for the original KORA metabolomics dataset. Due to different QC results across the two analytical kits, two metabolites, specifically PC aa C38:1 in *v*1 and C16:2 in *v*2, were excluded (see Table S2). The final dataset comprised 102 metabolites with 614 individuals who have data at both timepoints. These samples were divided into training, validation, and test sets with a ratio of 64%, 16%, and 20% respectively, corresponding to 393, 98, and 123 samples. The data in 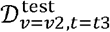 are masked for performance evaluation.

### Training the cGAN model

#### Architecture implementation

The cGAN^50^ extends the original GAN^51^ model by introducing additional information into the generation process, thereby providing more control over the generated output. In our context, the completion of missing views is conditioned on the data from observed views. Moreover, we enhanced the baseline cGAN model with an auxiliary classifier^28^ to ensure that the imputed view can be paired with the corresponding observed view.

Specifically, the generator *G*_cGAN_ learns the complex mappings between 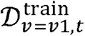 and 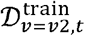, with *t*=*t*1 in our case. The reconstruction loss ℒ_*rec_cGAN*_ quantifies the differences between the actual and reconstructed data. The discriminator *D*_*cGAN*_ computes the adversarial loss ℒ_*adV_cGAN*_ by distinguishing if data are real 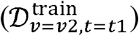 or generated 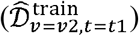. *D*_*cGAN*_ also has an auxiliary classifier that computes the auxiliary loss ℒ_*aux_cGAN*_ by predicting whether the pairs of views are real, i.e. 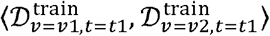, or fake, i.e.,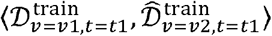. The final loss is defined as:

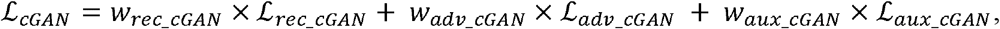

where *w*_*rec_cGAN*_, *w*_*adV_cGAN*_ and *w*_aux_cGAN_ represent weights for each loss. After training, the generator *G*_*cGAN*_ is applied to 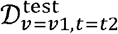 to generater 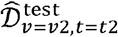.

The generator of the cGAN model consists of several residual blocks^52^ and uses the parametric rectified linear unit (PReLU)^53^ as its activation function. Each residual block includes batch normalization^54^ as necessary. MSE loss serves as ℒ_*rec_cGAN*_. Both MSE loss and binary cross-entropy (BCE) loss are considered for ℒ_*adV_cGAN*_ and ℒ_*aux_cGAN*_, determined by the hyperparameter tuning experiments. MSE and BCE loss are defined as:

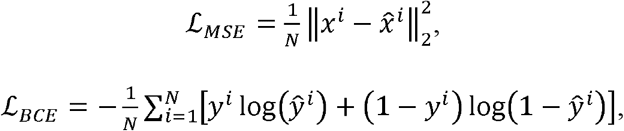

where *N* is the number of samples, *x*^*i*^ and 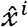 represents the true and estimated values for the *i*-th sample, while *y*^*i*^ and 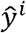 represents the true and predicted labels for the -the sample.

The cGAN model is trained with the Adam optimizer^55^, with a mini-batch size of 16 for the MGH COVID dataset and 32 for the two KORA-derived datasets. The model is implemented using PyTorch^56^ (v1.11.0), PyTorch Lightning^57^ (v1.6.4), and tensorboard^58^ (v2.10.0), and run on a graphics processing unit (GPU) operating compute unified device architecture (CUDA, v11.3.1).

#### Hyperparameter optimization

We utilized the training (64%) and validation (16%) sets from the KORA metabolomics dataset, which has the largest sample size among three benchmark datasets, to optimize the hyperparameters. We employed a grid search over various combinations of hidden layer size, numbers of residual block, batch normalization, and weights for different losses. We determined the number of training epochs based on early stopping triggered by the MSE reconstruction accuracy calculated on the observed data from the validation sets. Our experiments included variations in the number of hidden neurons for both the generator and discriminator, with options including 32, 64, 128, and 256. The number of residual blocks spanned from 2 to 6.

The final hyperparameters comprised five residual blocks of 64 neurons each for the generator and three hidden layers of 128 neurons each for the discriminator. Batch normalization was incorporated into the first four residual blocks of the generator and the last two layers of the discriminator to stabilize the learning process and accelerate convergence. A weight of 0.5 to ℒ_*rec_cGAN*_ and a weight of 0.25 to ℒ_*adV_cGAN*_ and ℒ_*aux_cGAN*_ were determined. MSE loss was selected for both ℒ_*adV_cGAN*_ and ℒ_*aux_cGAN*_. The determined hyperparameters were fixed and used in all evaluations.

### Training the LEOPARD

#### Architecture implementation

View-specific pre-layers 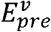 are used to embed input data 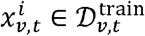 of different views into dimensionally uniform embeddings 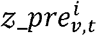. Here, *i* represents the data from the *i*-th individual. The representation disentangler of LEOPARD comprises a content encoder *E*_*c*_ and a temporal encoder *E*_*t*_, both shared by input data across different views and timepoints. This module learns a timepoint-invariant content representation 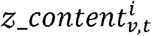 and temporal feature 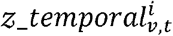 from 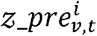. Following the encoding process, the generator *G* employs the AdaIN technique to re-entangle content representation and temporal knowledge:

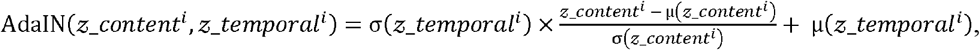

where μ and σ denote the mean and standard deviation operations respectively. View-specific post-layers 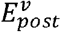 convert the re-entangled embeddings back to omics data 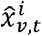. The discriminator *D* is trained to classify whether an input is a real sample or a generated output coming from *G* and 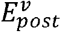. *D* produces the same number of outputs as the source classes of observed data, each corresponding to one view at one timepoint. For a sample belonging to source class *c*_*V,t*_, we penalize *D* during the update cycle of *D* if its output incorrectly the classifies a real data instance as false or a generated data instance as true for *c*_*V,t*_; when updating *G*, we only penalize *G* if *D* correctly identifies the generated data instance as false for *c*_*V,t*_

In our study, we defined the contrastive loss ℒ_*con*_ as the mean of the NT-Xent losses calculated separately for content and temporal representations. The NT-Xent loss is formulated as follows:

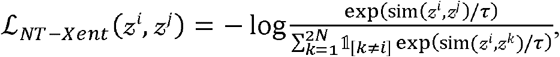

where *z*^*i*^ and *z*^*j*^ are embeddings of a positive pair (*i, j*), *τ* is a temperature factor that scales the similarities, and 𝟙_[*k* ≠ *i*]_ is an indicator function that equals 1 when *k* ≠ *i* and 0 otherwise. And sim (·)denotes cosine similarity, defined as:

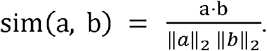

In each training iteration, we generate data for the missing view before calculating the loss. This allows the generated data to be factorized into corresponding content and temporal representations, further facilitating loss minimization. For the ℒ_*NT-Xent*_ calculated on content representations, positive pairs are defined as 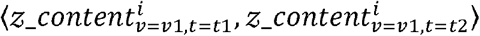 and 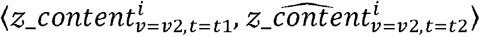, which are the same kind of content embeddings from the same individuals across different timepoints. Similarly, the positive pairs of temporal representations are 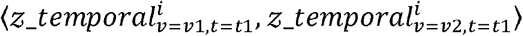 and 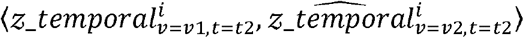. The representation loss ℒ _*rep*_ is the mean of the MSE losses calculated for content and temporal representations. For each type of representation, LEOPARD measures the MSE between the representation factorized from the actual data and reconstructed data. The reconstruction loss ℒ_*rec*_ quantifies the discrepancies between the actual and reconstructed data. Any missing values in the observed view are encoded as the mean values across each specific variable, and these mean-encoded values are excluded from the computation of ℒ_*rec*_ during back-propagation. This strategy enhances the robustness of LEOPARD in scenarios where input data contain missing values. The generator can arbitrarily produce data for any source classes given the content and temporal representations. To ensure the representation disentangler can capture the highly structured data pattern, we only compute the ℒ_*rec*_ on the data generated from content and temporal representations derived from different source classes. For instance, 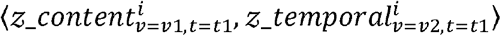 or 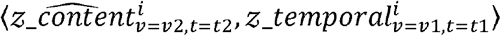.Data generated from representation pairs of the same views and timepoints, such as 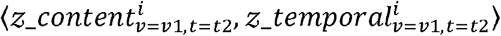 are not used for optimization. This design imposes additional restraints, and LEOPARD is tamed to learn more generalized representations, which helps prevent overfitting. Similar to the cGAN model described in the previous section, the adversarial loss ℒ_*adV*_ is also computed based on MSE. The final loss of LEOPARD is defined as:

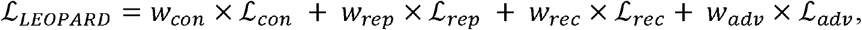

where *w*_*con*,_ ℒ_*con*,_ *w*_*rec*_ and *w*_*adv*_ the weights of the losses.

The encoders, generator, and discriminator of LEOPARD are built from blocks of layers without skip connections. Each block starts with a dense layer. An instance normalization layer^59^ is added after the dense layer for the content encoder. The encoders and generator use the PReLU as its activation function, while the discriminator uses the sigmoid function. A dropout layer^60^ is incorporated after the activation layer, where necessary.

The LEOPARD model is trained with the Adam optimizer, with a mini-batch size of 64. The model is implemented under the same computational environment as the cGAN model.

#### Ablation test

We conducted a comprehensive ablation study to assess the individual contributions of the four distinct losses incorporated into our LEOPARD architecture. By excluding each loss, we benchmarked the performance against a baseline setting that only utilizes reconstruction loss. The ablation test was performed with the training and validation sets from the KORA metabolomics dataset. Grid search was used to determine the optimal weights for the losses, and the median of PB computed from the validation set was used to quantify the performance. The network layer numbers and sizes were consistent during the evaluation. We used three hidden layers for generator and encoders, with each layer containing 64 neurons. The weight for reconstruction loss was fixed at 1, while the weights for the other three losses varied across 0.01, 0.05, 0.1, 0.5, and 1. The number of training epochs was determined by the model’s saturation point in learning, which is when the median of PB computed on the validation set ceased to decrease significantly. Our experiments show include *w*_*rec*_ =1, *w*_*con*_ =0.1, *w*_*rep*_ =0.1, and *w*_*adv*_ =1,. The performance of each loss that all four losses contribute to lowering PB in the imputed data. The optimal weights combination at their optimal weights is summarized in Extended Fig. 4.

#### Further hyperparameter optimization

The training and validation sets from the KORA metabolomics dataset were further used to optimize LEOPARD’s hyperparameters, aiming to effectively capture data structure for existing data reconstruction and achieve robust generalization for missing data imputation. The weights for different losses have been determined in the ablation test. We then conducted a grid search across various combinations of hidden layer size, hidden layer number, dropout rate, projection head and temperature for contrastive loss.

The number of hidden neurons within the encoders, generator, and discriminator varied across 32, 64, 128, and 256, with the number of layers ranging from 2 to 4, and dropout rates of 0%, 30%, and 50%. Our findings show that higher numbers of hidden neurons and layers tended to yield worse performance in terms of median PB (Extended Fig. 5). LEOPARD was configured with three 64-neuron layers incorporated into both content and temporal encoders and the generator. The discriminator included two hidden layers, each having 128 neurons. Dropout was not used.

Projection head and temperature are two important hyperparameters that control the performance of contrastive learning. The projection head is a compact network consisting of one full connected hidden layer with the same layer size as the input dimension, a rectified linear unit (ReLU)^61^ and one output layer. The temperature is a scalar that scales the similarities before the softmax operation. Some previous experiments performed on image datasets emphasized the importance of the projection head and reported different output sizes yielded similar results^29^. We evaluated the performance of LEOPARD both without a projection head and with a projection head of the output size varying across 16, 32, 64, 128, 256, and 512. The temperature is fined-tuned across 0.05, 0.1, 0.5, 1, 5, 10, and 30. Based on our experiments, LEOPARD is trained with a temperature of 0.05 and without using a projection head (Extended Fig. 6).

The determined hyperparameters, including loss weights, remained unchanged in all our performance evaluations.

### Representation disentanglement

The disentanglement of content and temporal representations was evaluated using the KORA multi-omics dataset. LEOPARD was trained for 600 epochs, for each of which the disentanglement progress was visualized with the following steps: First, content and temporal representations were factorized from the metabolomics (S4 and F4) and proteomics data (S4). Then the generator imputed the proteomics data (F4) by incorporating the temporal information from the metabolomics data (F4) into the content representation from the proteomics data (S4). The generated proteomic data (F4) were then fed to the content and temporal encoders to extract the corresponding representations. Subsequently, these content or temporal representations of both the observed and imputed data were standardized to ensure all latent variables had a mean of zero and a standard deviation of one. Afterward, two separate UMAP models were built using the R package umap^62^, each fitted to the content and temporal representations, with a configuration of *n_neighbors* = 15 and *min_dist* = 0.1. Lastly, scatter plots were generated using the R packages ggplot2^63^ and ggsci^64^. Each point in the plot represents an individual sample, and the color indicates the data sources. The visualization epochs were selected experimentally to illustrate the progress of representation disentanglement during the training process.

### Performance evaluation

The LEOPARD and cGAN models were trained using the hyperparameters previously described. For missForest, the imputation was performed using a 100-tree random forest^65^ model, with the maximum number of iterations (*maxit*) set to 10. Multiple imputations by PMM, LM, and GLMM were performed using the R packages mice^35^ and micemd^66^. Each method’s imputations were performed five times (*m* = 5) with a *maxit* value set to five. The PMM model was built using argument *method* = “pmm”. When *obsNum* = 0, data of *v*2 at *t*2 are assumed to be completely missing. In this scenario, the LM method was trained using *method* = “norm”. When *obsNum* is a non-zero value, the GLMM model was built using *method* = “2l.glm.norm”.

All methods used only the data in the training sets to build imputation models. Their performance was evaluated on 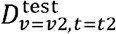. Different imputation methods may require specific data structures for input: cGAN and LEOPARD first build imputation models using training data, then apply the built models to test set to complete missing views. In contrast, the input data for other methods can be an incomplete matrix with missing values coded as NA. We adapted the input data accordingly to accommodate these specific requirements:

- Method cGAN only learns from samples where both views are present. Therefore, its training data only included training data from the first timepoint (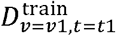 and 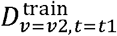) and data of different *obsNum* from the second timepoint (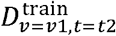 and 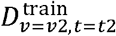).
- LEOPARD can additionally learn from data where only one view is available. In addition to 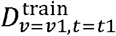 and 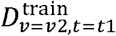, the entire 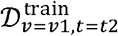 was included in its training. The variation of *obsNum* only affected the number of observed samples from 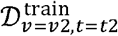.
- The input data for missForest combined training data (including 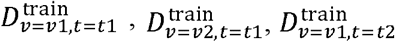, and data of different *obsNum* from 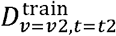) and test data 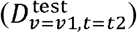, with NA filling the masked data in the matrix.
- For the multiple imputation methods in MICE family, the input data were constructed with training data (identical to that used for missForest) and test data (including 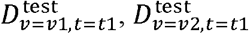 and 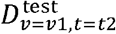). To ensure test data remained unused for model training, a logical vector with TRUE assigned to test samples was passed to the *ignore* argument. Masked values were filled with NA in the matrix.
- When building the GLMM model, the input data additionally contained sample IDs and timepoint labels. A constant residual error variance is assumed for all individuals. Building GLMM model for a large dataset is extremely time consuming; thus, for the MGH COVID dataset, we selected the top 100 highly Spearman-correlated proteins for each protein requiring imputation. The selected proteins were incorporated into the imputation process by passing to the argument *predictorMatrix*.

PB was selected as performance metric as it quantifies the relative deviation of imputed values from actual observations, offering a more straightforward interpretation compared to metrics like RMSE and MAE. PB was calculated for each variable using the formula:

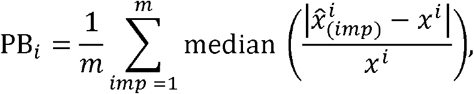

where 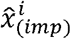 is the imputed value for the *i*-th variable from the *imp*-th imputation, while *m* is the number of imputations. For single imputation methods, LEOPARD, cGAN, and missForest, *m* = 1. PB results for each imputation method were visualized using dot and box plots, with each dot representing a variable in the specific dataset.

For the evaluation using UMAP, we first fitted a UMAP model using the data of 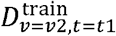 And 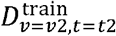. We then used the fitted model to embed the data of 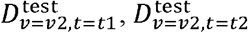, and 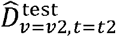, where 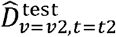 represents the imputed data for 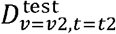 produced by different imputation methods. For PMM, LM, and GLMM, 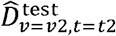 is the average of all estimates from their multiple imputations. An imputation method is considered effective if the distribution of 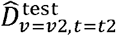 embeddings is highly similar to that of the 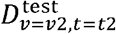 embeddings. The UMAP models were fitted with the identical configurations described in the prior section.

### Regression analyses

We used observed and imputed test set data from the KORA metabolomics and multi-omics datasets for two regression analyses. We employed multivariate linear regression models for each of the observed or imputed data. The imputed data of LEOPARD, cGAN, missForest, PMM, and LM were obtained under the setting of *obsNum* = 0.

For the KORA metabolomics dataset, we used the concentration of each metabolite as the response variable and age as the predictor variable, while controlling for sex, to detect age-associated metabolites. For the KORA multi-omics dataset, we used NPX values of each protein as the response and eGFR as the predictor, controlling for sex and age, to detect eGFR-associated proteins. In both analyses, we applied a Bonferroni correction to adjust the *P*-value significance threshold to mitigate the risk of false positives in multiple testing.

### Classification analyses

We also used observed and imputed test set data from the KORA metabolomics and multi-omics dataset for CKD prediction. CKD cases were determined based on their eGFR values, which were computed from serum creatinine, sex, race, and age, using the Chronic Kidney Disease Epidemiology Collaboration (CKD-EPI) equation^67^. The data imputed by LEOPARD, cGAN, missForest, PMM, and LM were obtained under the setting of *obsNum* = 0.

For each type of raw or imputed data, we trained a BRF model using the Python library imbalanced-learn^68^. This model was specifically selected to address the dataset imbalance and reduce the risk of overfitting to the majority class. All models were trained with default hyperparameters (*criterion* = “gini”, *min_samples_split* = 2, *min_samples_leaf* = 1, *max_features* = “sqrt”, *bootstrap* = True), except for *n_estimators* = 1000 and *class_weight* = “balanced_subsample”. Due to the limited sample size, we validated the performance using the LOOCV strategy, allowing maximal use of data for both training and validation. Performance metrics were calculated using the R package caret^69^. These metrics provided a comprehensive understanding of the predictive power of the observed and imputed data.

The ROC curves were plotted to illustrate the trade-off between sensitivity and 1-specificity at varying decision thresholds. Considering the imbalance in our dataset, and with our primary interest in the positive class, which is also the minority, we further plotted PR curves to depict the trade-off between precision and recall at different thresholds for the classifiers trained with the different data. For PR curves, the baseline performance of a non-discriminative model was determined by the proportion of positive cases (56/416 = 0.135 for the KORA metabolomics dataset and 36/212 = 0.170 for the KORA multi-omics dataset). Both the ROC and PR curves were plotted using the R package precrec^70^.

### Arbitrary temporal knowledge transfer

In the previous evaluation, we assessed the performance of each method on 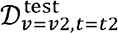 from the benchmark datasets. We then extended our analysis by evaluating LEOPARD’s performance on individually masked test sets: 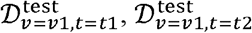, and 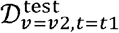 from KORA metabolomics dataset. This approach allows us to assess LEOPARD’s capability to arbitrarily complete any views at any timepoints within this dataset. Moreover, LEOPARD was evaluated using the expanded KORA metabolomics dataset to complete masked test set 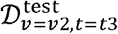, using the training data from 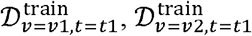, and 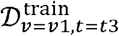. This analysis further enables us to explore LEOPARD’s capability to complete views across a long time span. LEOPARD was trained using the same hyperparameters as we used in the previous experiments.

## Data availability

The MGH COVID data, published by the original authors, have been deposited to Mendeley Data (http://dx.doi.org/10.17632/nf853r8xsj). The KORA data are governed by the General Data Protection Regulation (GDPR) and national data protection laws, with additional restrictions imposed by the Ethics Committee of the Bavarian Chamber of Physicians to ensure data privacy of the study participants. Therefore, the data cannot be made freely available in a public repository. However, researchers with a legitimate interest in accessing the data may submit a request through an individual project agreement with KORA via the online portal (https://www.helmholtz-munich.de/en/epi/cohort/kora). Upon receipt of the request, the data access committee will review the application and, subject to approval, provide the researcher with a data usage agreement. The specific variable names and sample IDs for each dataset, determined following the QC process, can be found in Table S1-S3.

## Code availability

The source code and implementation details of LEOPARD are freely available at our GitHub repository (https://github.com/HAN-Siyu/LEOPARD). Detailed documentation and examples can be found in the Manual, which is also available at this repository.

## Acknowledgements

This project has received funding from the Innovative Medicines Initiative 2 Joint Undertaking (JU) under grant agreement No 821508 (CARDIATEAM). The JU receives support from the European Union’s Horizon 2020 research and innovation programme and the European Federation of Pharmaceutical Industries and Associations (EFPIA).

The German Diabetes Center is supported by the German Federal Ministry of Health (Berlin, Germany) and the Ministry of Science and Culture in North-Rhine Westphalia (Düsseldorf, Germany). This study was supported in part by a grant from the German Federal Ministry of Education and Research to the German Center for Diabetes Research (DZD).

The KORA study was initiated and financed by the Helmholtz Zentrum Mu□nchen – German Research Center for Environmental Health, which is funded by the German Federal Ministry of Education and Research (BMBF) and by the State of Bavaria. Data collection in the KORA study is done in cooperation with the University Hospital of Augsburg.

We express our appreciation to all KORA study participants for their blood donation and time. We thank all participants for their long-term commitment to the KORA study, the staff for data collection and research data management and the members of the KORA Study Group (https://www.helmholtz-munich.de/en/epi/cohort/kora) who are responsible for the design and conduct of the KORA study.

We also extend our gratitude to all participants of the MGH COVID study, as well as the dedicated staff who have participated in the study’s design, data collection, and project management.

The authors are grateful to Prof. Dr. Barbara Thorand (Institute of Epidemiology, Helmholtz Zentrum München) for her contribution to the collection of KORA proteomics data and for her constructive suggestions to improve this study.

The authors thank Yunhsiu Tai, Ming Cheng, Ruoyu Wang, Yuan Guo, Linrui Fan for their supports and suggestions during the development of LEOPARD.

## Author contributions

Siyu Han designed and carried out the analyses, implemented the algorithms, interpreted the result, and wrote the manuscript.

Shixiang Yu, and Ying Li assisted in the development of the method.

Mengya Shi, Makoto Harada, Jianhong Ge, Flora Sam, Giuseppe Matullo assisted in interpreting the results.

Jiesheng Lin, Cornelia Prehn, Agnes Petrera, Jerzy Adamski, Karsten Suhre, Christian Gieger, Stefanie M. Hauck, Christian Herder, Michael Roden, Annette Peters, and Rui Wang-Sattler performed data acquisition and preparation of the KORA cohort data.

Na Cai and Francesco Paolo Casale assisted in the analysis and revised the manuscript.

Rui Wang-Sattler supervised the analyses, interpreted the results, and revised the manuscript. All authors reviewed the final manuscript.

## Additional information

**Correspondence and requests for materials** should be addressed to Rui Wang-Sattler.

**Extended Table 1.** Method performance of CKD prediction on KORA metabolomics dataset.

| Method | Sensitivity | Specificity | Precision | Balanced Accuracy | F1 Score | AUROC | PRRPC |
| --- | --- | --- | --- | --- | --- | --- | --- |
| *groundtruth* | 0.643 | 0.800 | 0.333 | 0.721 | 0.439 | 0.831 | 0.474 |
| PMM | 0.125 | **0.897** | 0.159 | 0.511 | 0.140 | 0.543 | 0.186 |
| LM | 0.161 | 0.903 | 0.205 | 0.532 | 0.180 | 0.590 | 0.166 |
| missForest | 0.268 | 0.744 | 0.140 | 0.506 | 0.184 | 0.486 | 0.135 |
| cGAN | 0.268 | 0.789 | 0.165 | 0.528 | 0.204 | 0.555 | 0.177 |
| LEOPARD | **0.571** | 0.747 | **0.260** | **0.659** | **0.358** | **0.719** | **0.268** |

Bold indicates the highest performance among imputation methods.

**Extended Table 2.** Method performance of CKD prediction on KORA multi-omics dataset.

| Method | Sensitivity | Specificity | Precision | Balanced Accuracy | F1 Score | AUROC | PRRPC |
| --- | --- | --- | --- | --- | --- | --- | --- |
| *groundtruth* | 0.778 | 0.778 | 0.418 | 0.778 | 0.544 | 0.862 | 0.593 |
| PMM | **0.667** | 0.494 | 0.212 | 0.580 | 0.322 | 0.631 | 0.282 |
| LM | 0.611 | 0.562 | 0.222 | 0.587 | 0.326 | 0.665 | 0.339 |
| missForest | 0.583 | 0.562 | 0.214 | 0.573 | 0.313 | 0.589 | 0.230 |
| cGAN | 0.500 | 0.545 | 0.184 | 0.523 | 0.269 | 0.551 | 0.198 |
| LEOPARD | **0.667** | **0.665** | **0.289** | **0.666** | **0.403** | **0.725** | **0.435** |

Bold indicates the highest performance among imputation methods.

**Extended Fig. 1.**
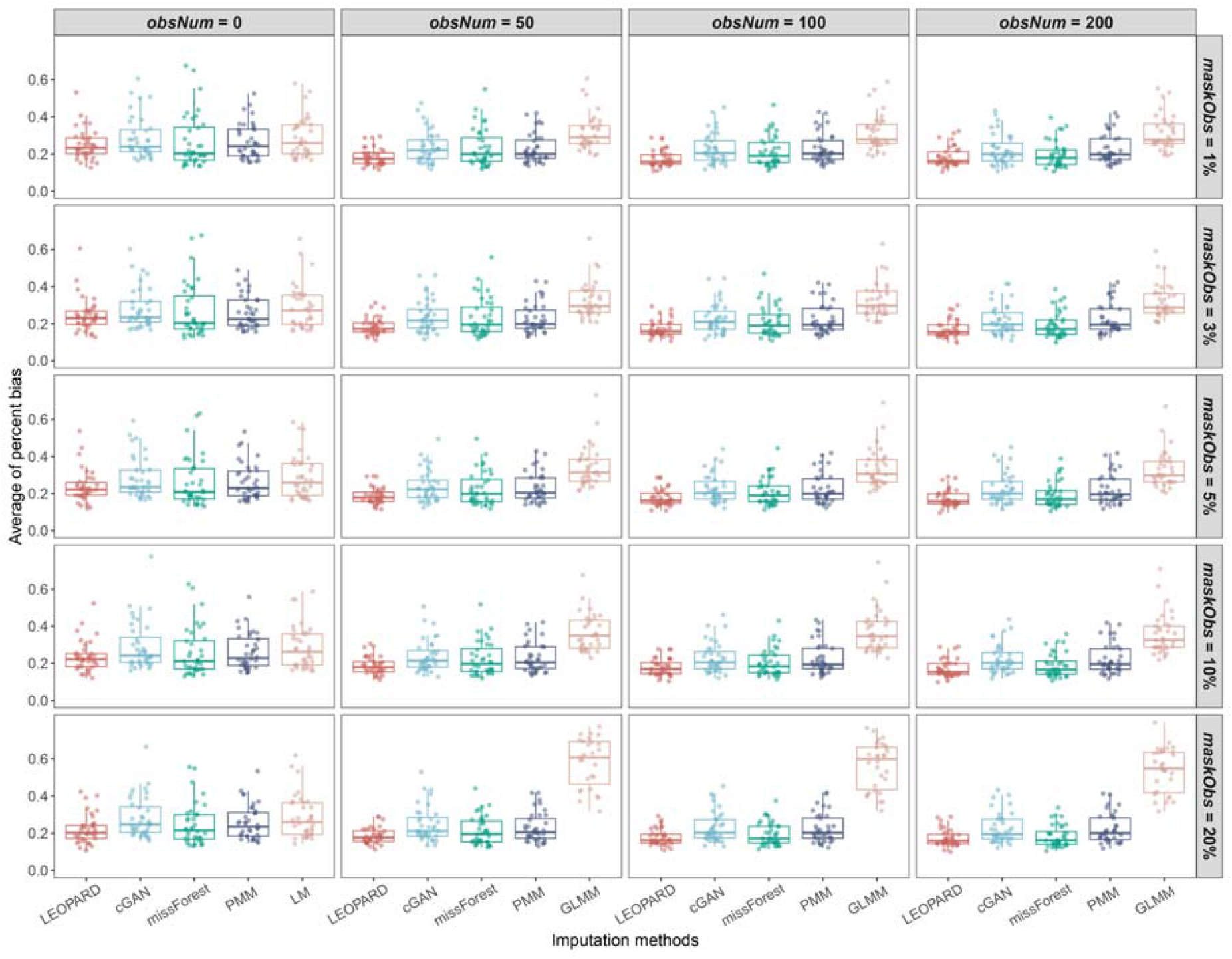
Performance of missing view completion when observed views have missing values. The average PB are computed for 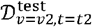 of the KORA metabolomics dataset across 10 repeated completions, under each proportion of masked data points in the observed views (*maskObs*). In each repetition, the data points are masked randomly. Each dot represents a PB value for a variable. Please note that LM is used instead of GLMM when *obsNum* = 0.

**Extended Fig. 2.**
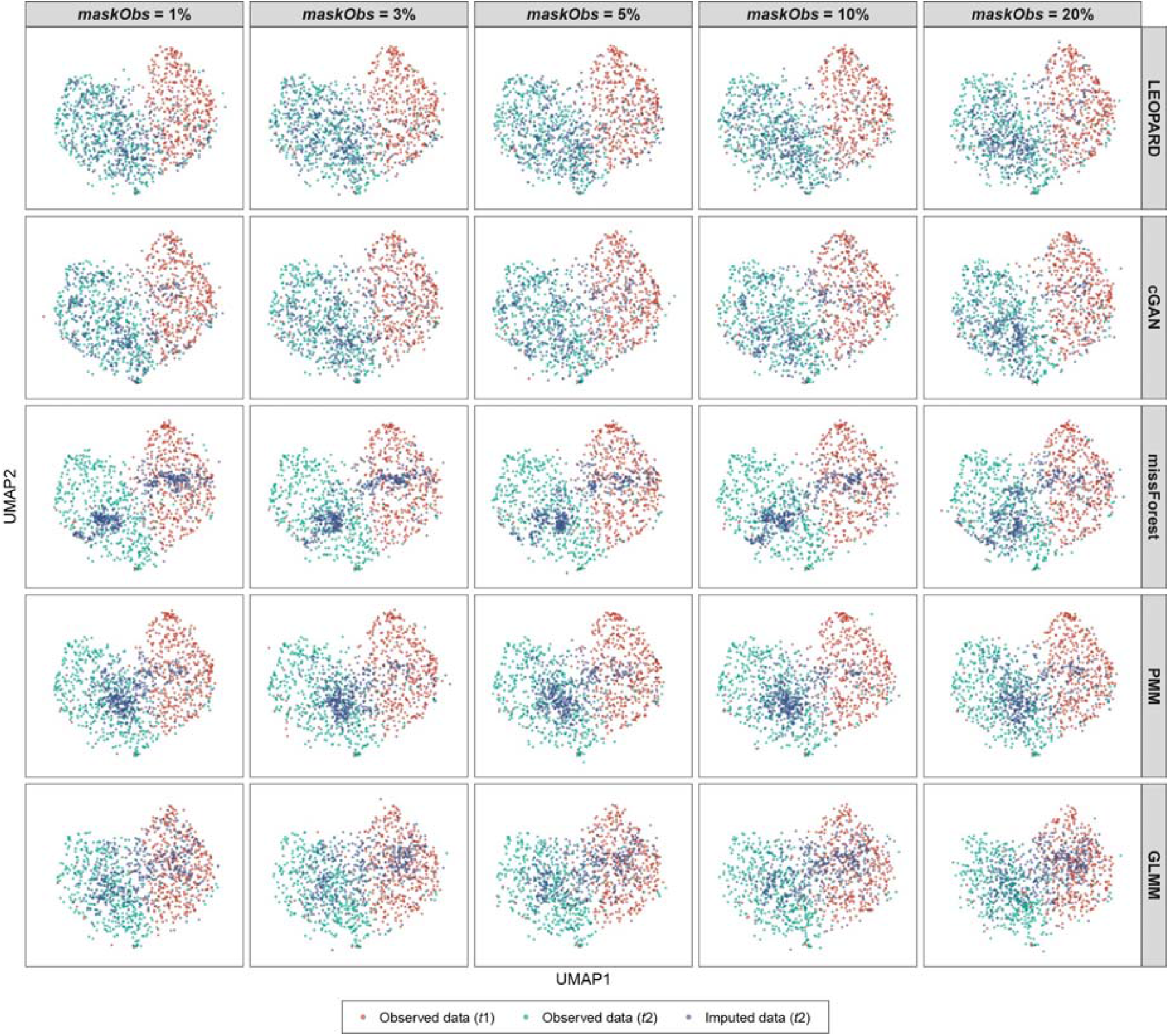
UMAP visualizes the imputed data obtained when observed views have missing values. UMAP models are initially fitted with the training data from the KORA multi-omics dataset (**c**, *t*1 : S4, *t*2 : F4). Subsequently, the trained models are applied to the corresponding observed data (represented by red and blue dots for *t*1 and *t*2) and the data imputed by different methods (represented by green dots) under *obsNum* = 200 and varying *maskObs*. Please note that, for each *maskObs*, only the repetition that exhibits the lowest median of PB are visualized. The distributions of red and blue dots illustrate the variation across the two timepoints, while the similarity between the distributions of blue and green dots indicates the quality of the imputed data. A high degree of similarity suggests a strong

**Extended Fig. 3.**
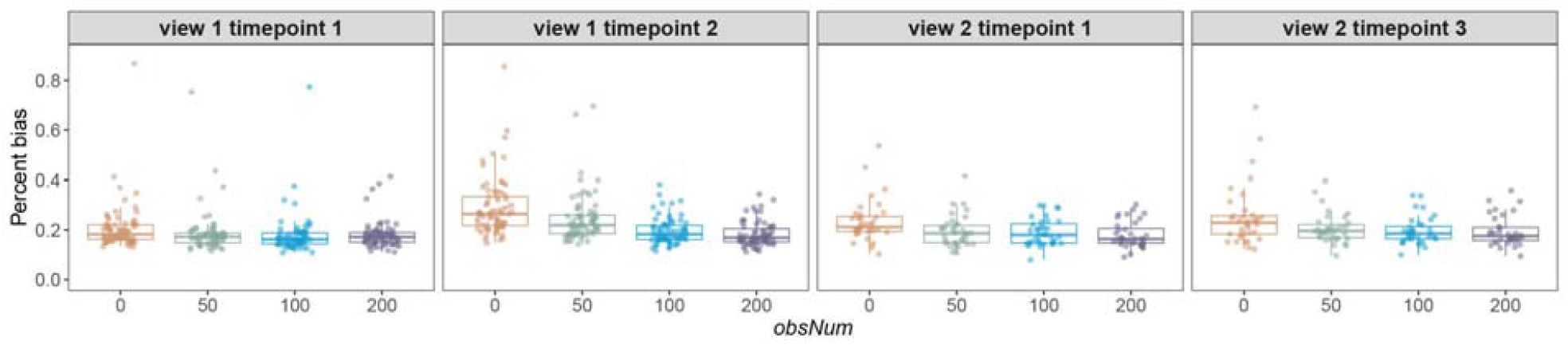
Performance of arbitrary temporal knowledge transfer. The first three panels illustrate LEOPARD’s performance evaluated on 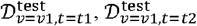, and 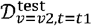of the KORA metabolomics dataset, with varying *obsNum*. The fourth panel displays LEOPARD’s performance on the extended KORA metabolomics dataset, where it completes the missing view 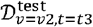 using the training data at *t*1. Each dot represents a PB value for a variable.

**Extended Fig. 4.**
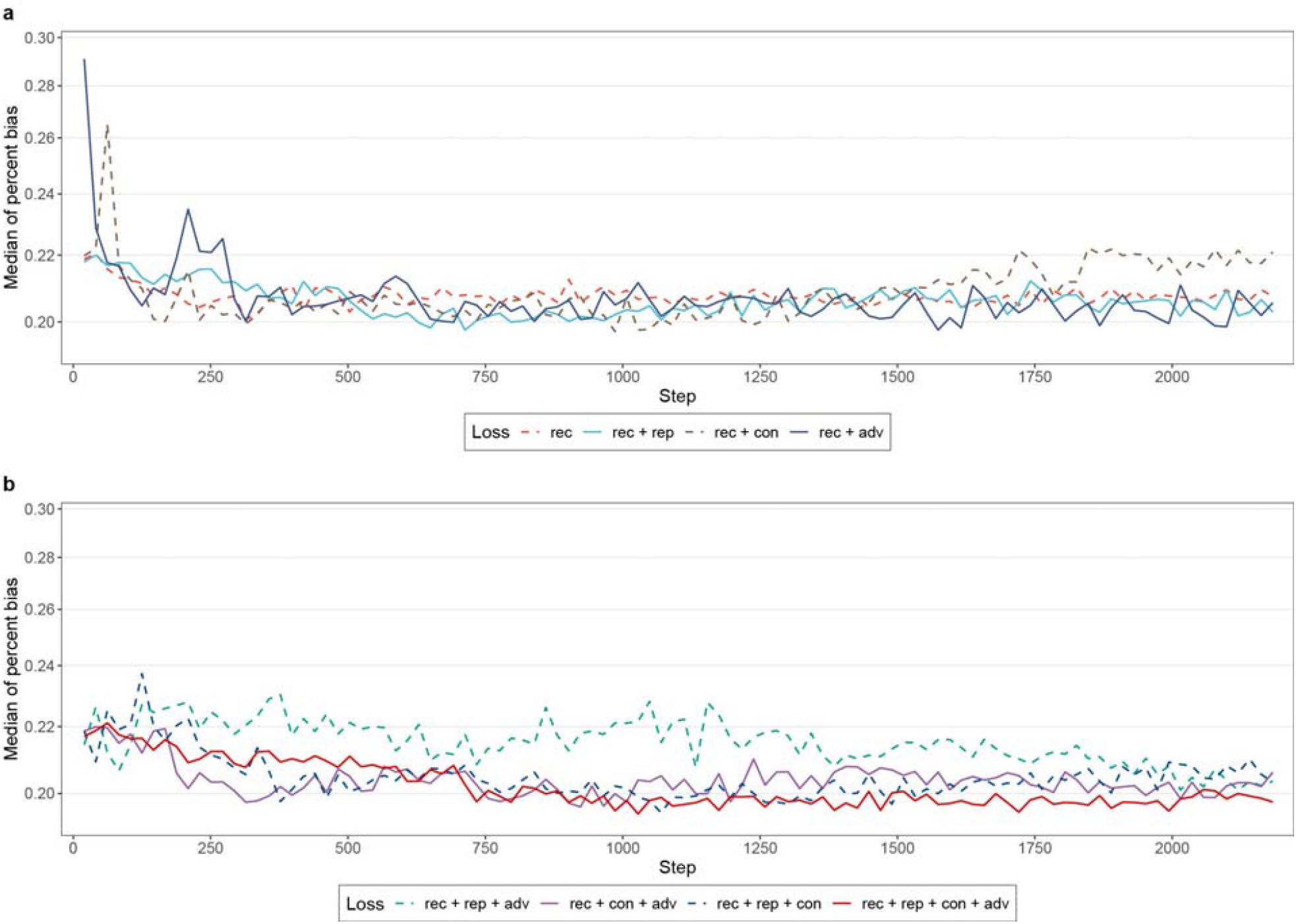
Ablation test evaluates the contribution of different losses. The ablation test evaluates the contribution of four losses: reconstructive loss (rec), representation loss (rep), contrastive loss (con), and adversarial loss (adv). Grid search strategy is used to determine the optimal weights for different losses, and median PB computed from the validation set is used to quantify the performance. The network layers are consistent during the evaluation. The plot shows the performance of reconstruction loss combined with (**a**) a single loss and (**b**) multiple losses, using the optimal weight of each loss. The weight for rec is fixed at 1, and the weights for the other losses vary across 0.01, 0.05, 0.1, 0.5, 1. The optimal weights for different losses in each combination are: rec + rep (*W*_*rep*_ = 0.01), rec + con (*W*_*con*_ = 0.01), rec + adv (*W*_*adv*_ = 0.5), rec + rep (*W*_*rep*_ = 0.01) + adv (*W*_*adv*_ = 1), rec + con (*W*_*con*_ = 0.01) + adv (*W*_*adv*_ = 1), rec + rep (*W*_*rep*_ = 0.5) + con (*W*_*con*_ = 0.01), rec + rep (*W*_*rep*_ = 0.1) + con (*W*_*con*_ = 0.1) + adv (*W*_*adv*_ = 1).

**Extended Fig. 5.**
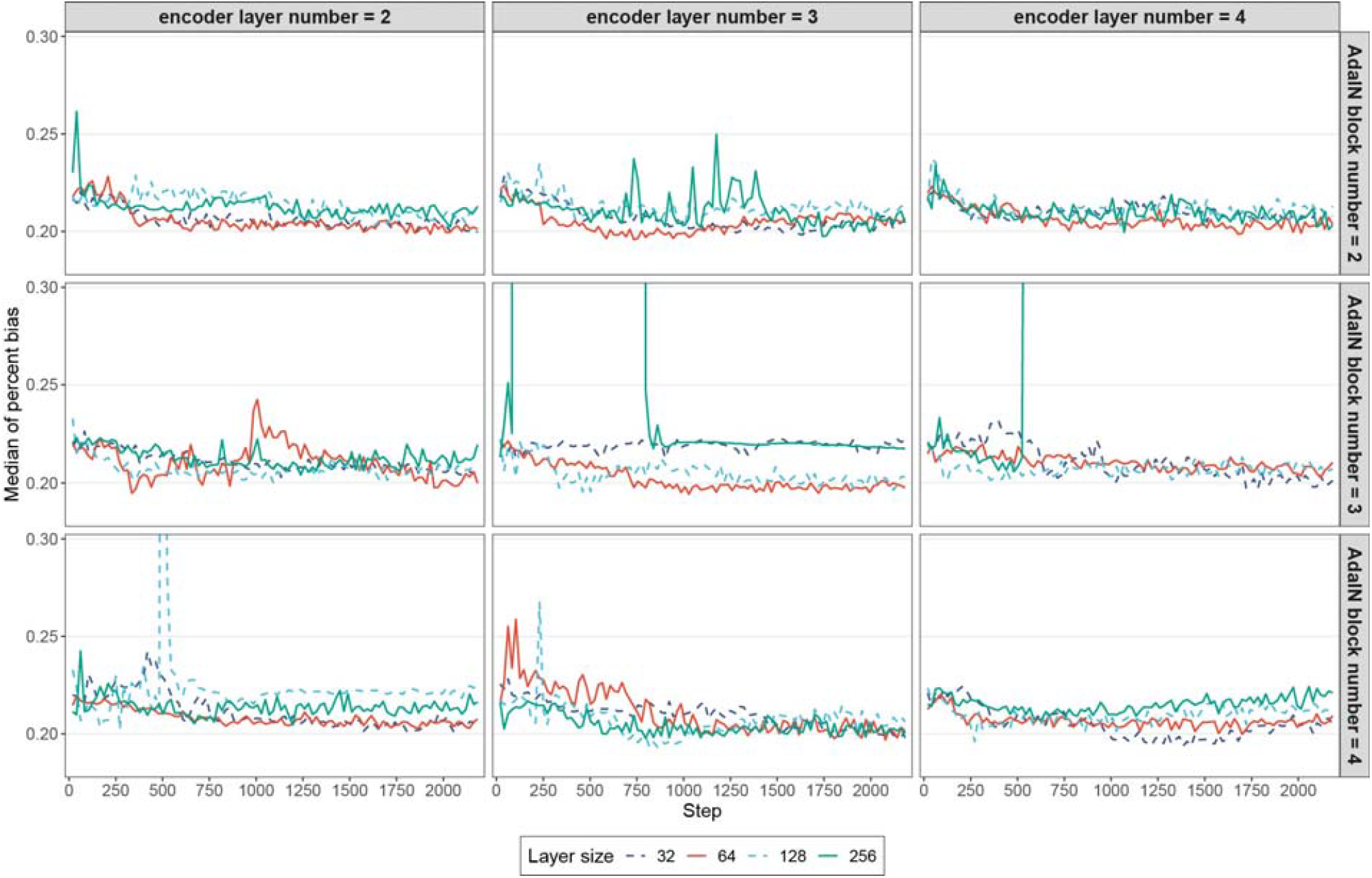
Hyperparameter tuning on network width and depth. Grid search strategy is used to determine the optimal layer number for the content and temporal encoders, AdaIN block number for the generator, and size for each layer. Median PB computed from the validation set is used to quantify the performance. The model achieves the lowest median PB with a setting of three layers for the encoders, each containing 64 neurons, and three AdaIN blocks for the generators, each including a 64-neuron layer.

**Extended Fig. 6.**
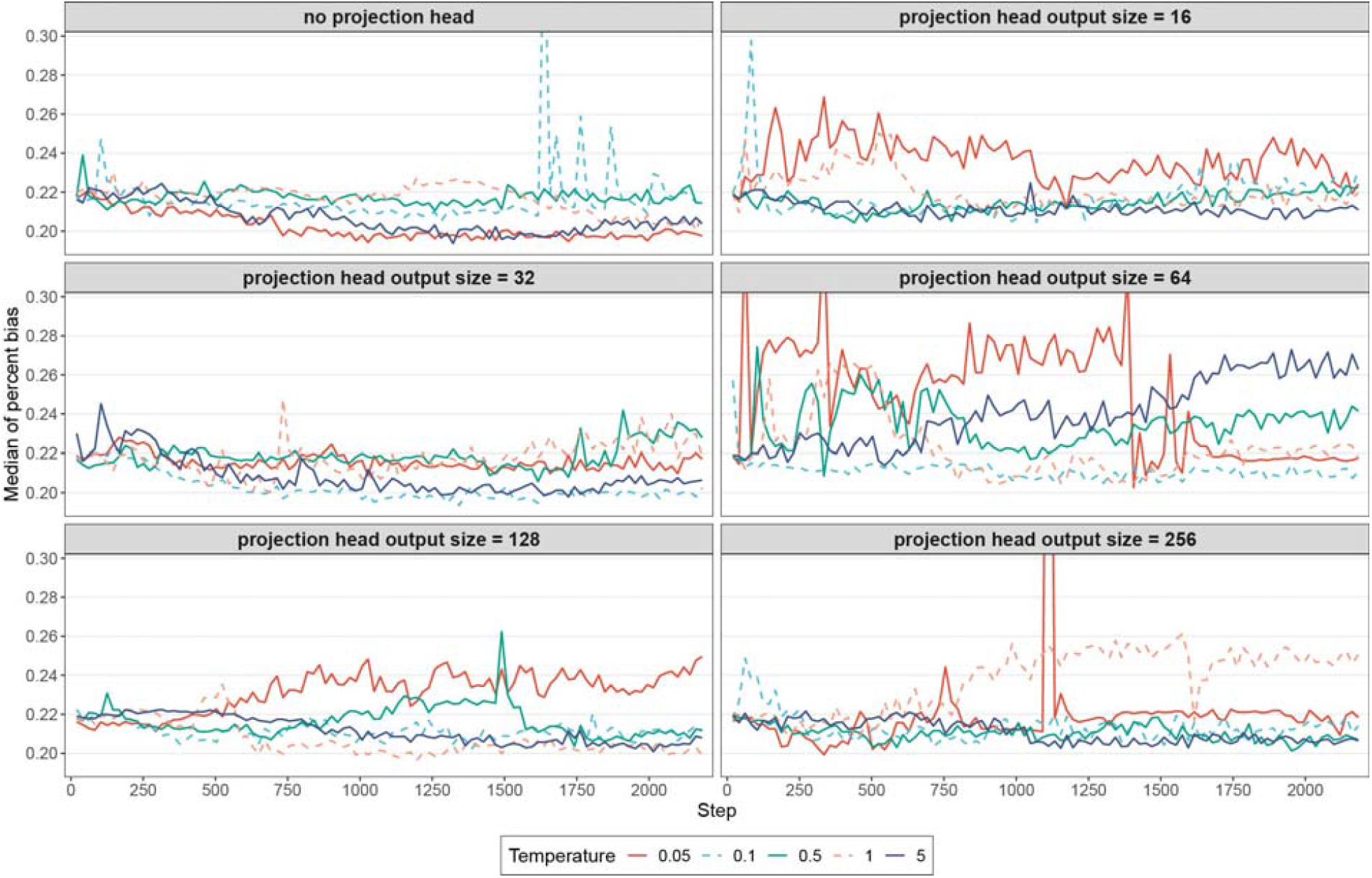
Hyperparameter tuning on projection head and temperature in contrastive loss. Based on the hyperparameters determined in the previous experiments, we further fine-tuned projection head and temperature used in our contrastive loss. We use grid search to evaluate the performance of LEOPARD without a projection head and with a projection head of different output sizes. The temperature varies across 0.05, 0.1, 0.5, 1, 5, 10, and 30. Based on our experiments, LEOPARD is trained with a temperature of 0.05 and without using a projection head.

