## Supplementary Figures S1-S5 for "LEOPARD: missing view completion for multi-timepoint omics data via representation disentanglement and temporal knowledge transfer"

**This document contains figures that evaluate each individual imputation performed by the multiple imputation methods assessed in our study.**

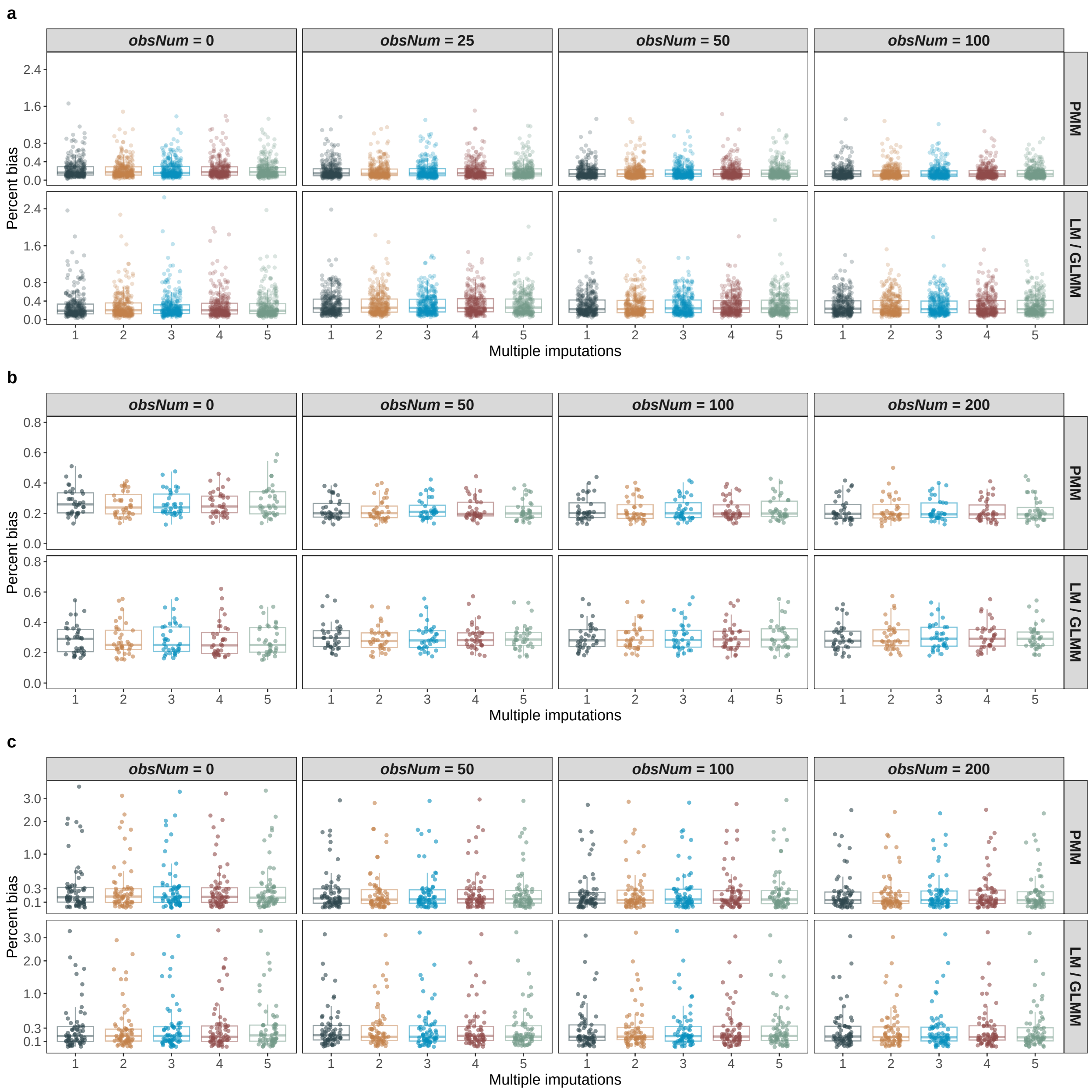

**Fig. S1** PB of each individual imputation of methods PMM and LM/GLMM calculated on $\mathcal{D}^{\text{test}}_{v=v2,t=t2}$ of the MGH COVID proteomics dataset (**a**), KORA metabolomics dataset (**b**), and KORA multi-omics dataset (**c**). **a**-**c**, with varying $obsNum$. Each dot represents a PB value for a variable. Please note that LM is used for imputation instead of GLMM when $obsNum = 0$.

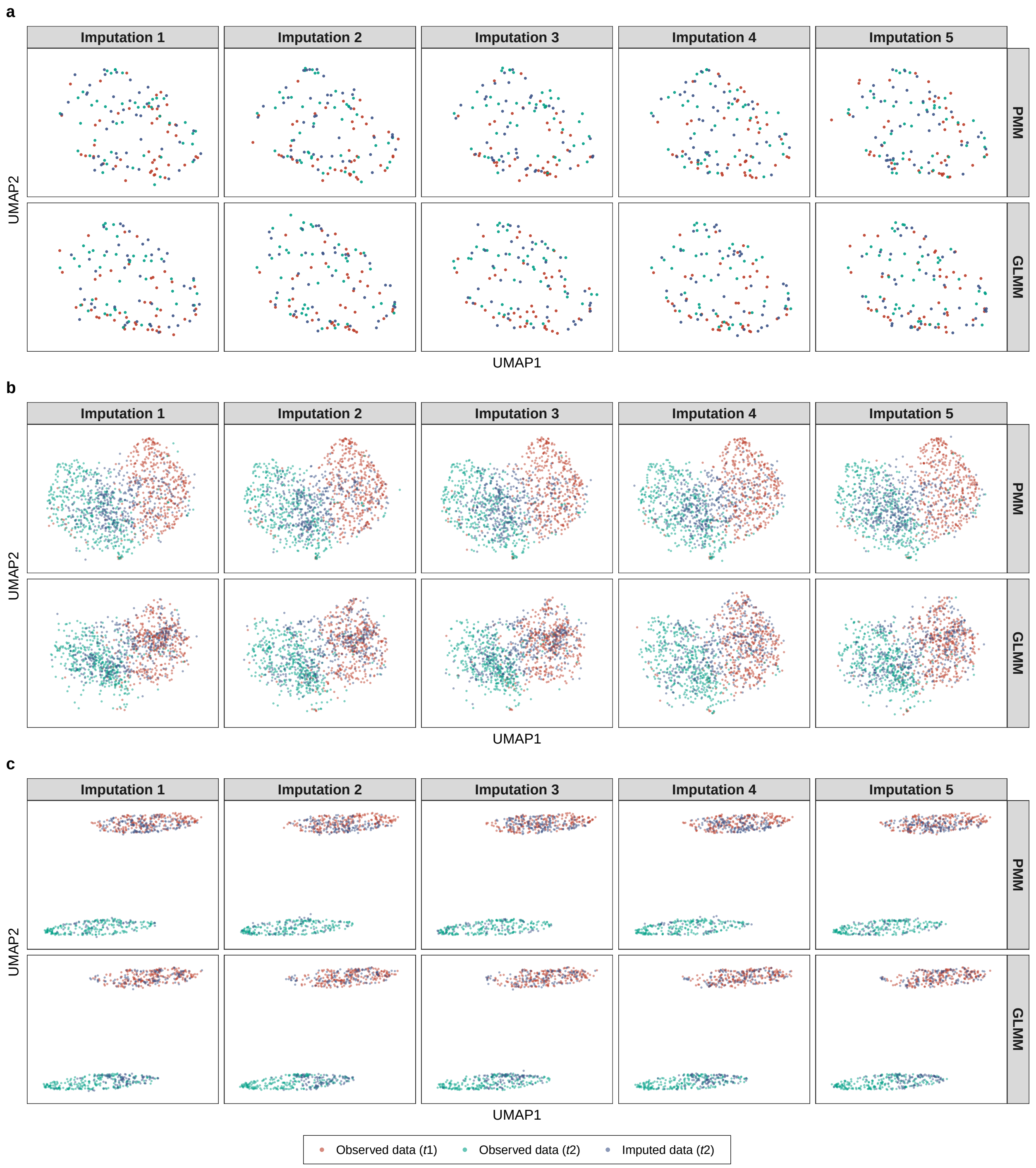

**Fig. S2** UMAP representations of each individual imputation of methods PMM and GLMM and corresponding observed data of three benchmark datasets. **a**-**c**, UMAP models are initially fitted with the training data from the MGH COVID proteomics dataset (**a**, $t1$: D0, $t2$: D3), KORA metabolomics dataset (**b**, $t1$: F4, $t2$: FF4), and KORA multi-omics dataset (**c**, $t1$: S4, $t2$: F4). Subsequently, the trained models are applied to the corresponding observed data (represented by red and blue dots for $t1$ and $t2$) and each individual imputation of PMM and GLMM (represented by green dots) under the setting of $obsNum = 100$ for the MGH COVID dataset and $obsNum = 200$ for the two KORA-derived datasets. The distributions of red and blue dots illustrate the variation across the two timepoints, while the similarity between the distributions of blue and green dots indicates the quality of the imputed data. A high degree of similarity suggests a strong resemblance between the imputed and observed data.

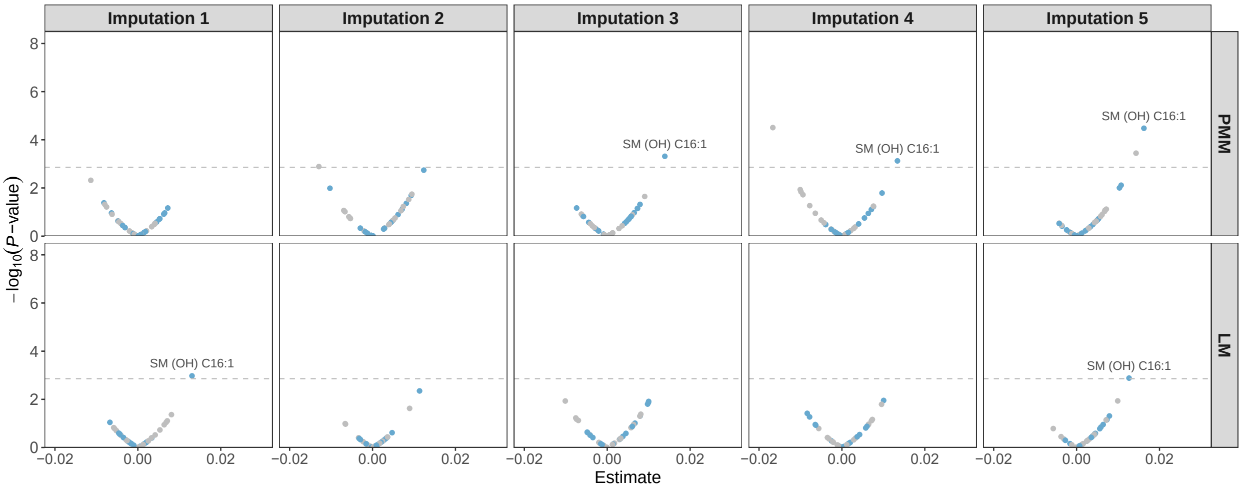

**Fig. S3** The age-associate metabolites identified from each individual imputation of methods PMM and LM. The evaluation is performed on $\mathcal{D}^{\text{test}}_{v=v2,t=t2}$ ($N = 417$) of the KORA metabolomics dataset, under $obsNum = 0$. 18 significant metabolites ($P < 0.05/36$) identified from the observed data are shown in blue. Replicated metabolites from the imputed data ($obsNum = 0$) are marked with labels.

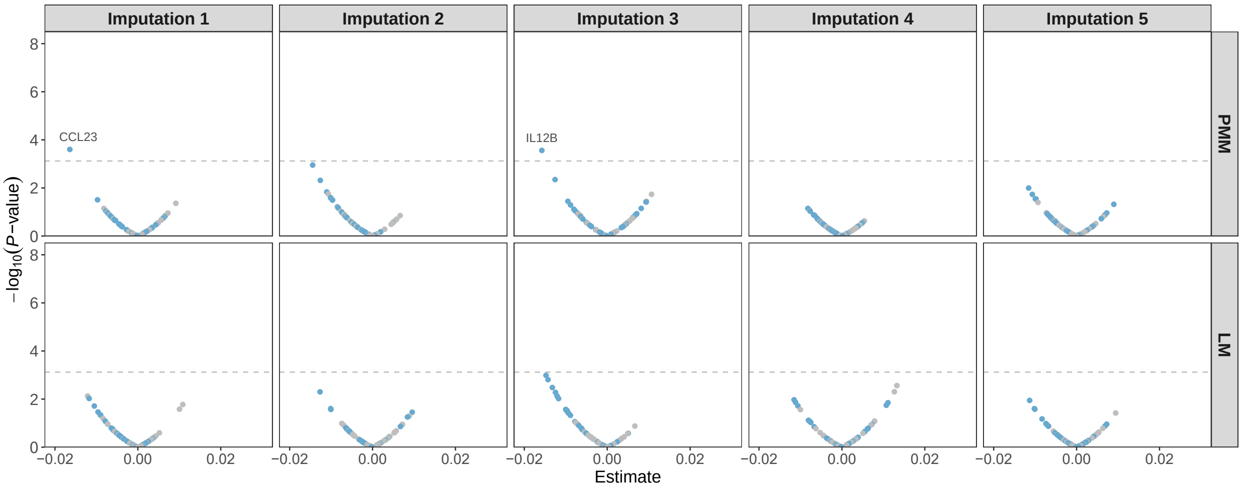

**Fig. S4** The eGFR-associated proteins identified from each individual imputation of methods PMM and LM. The evaluation is performed on $\mathcal{D}^{\text{test}}_{v=v2,t=t2}$ ($N = 212$) of the KORA multi-omics dataset, under $obsNum = 0$. 28 significant metabolites ($P < 0.05/66$) identified from the observed data are shown in blue. Replicated metabolites from the imputed data ($obsNum = 0$) are marked with labels.

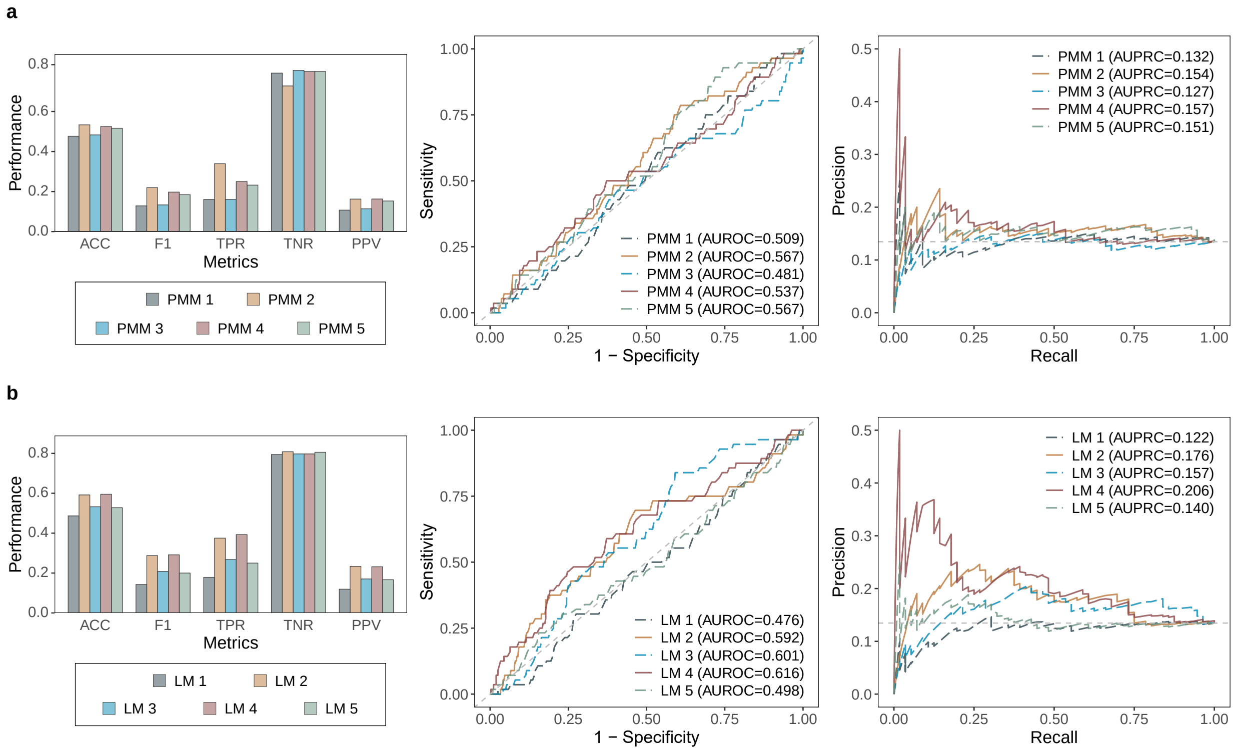

**Fig. S5** The performance of CKD prediction on the KORA metabolomics dataset using each individual imputation of methods PMM (**a**) and LM (**b**). Methods are evaluated on $\mathcal{D}^{\text{test}}_{v=v2,t=t2}$ ($N = 416$, $N_{positive} = 56$, $N_{negative} = 360$), under $obsNum = 0$. Models are trained using the BRF algorithm with identical hyperparameters and evaluated using LOOCV. The barplot (left) shows multi-metric performance. The dashed lines in the ROC (middle) and PR (right) curves represent the performance of a hypothetical model with no predictive capability.

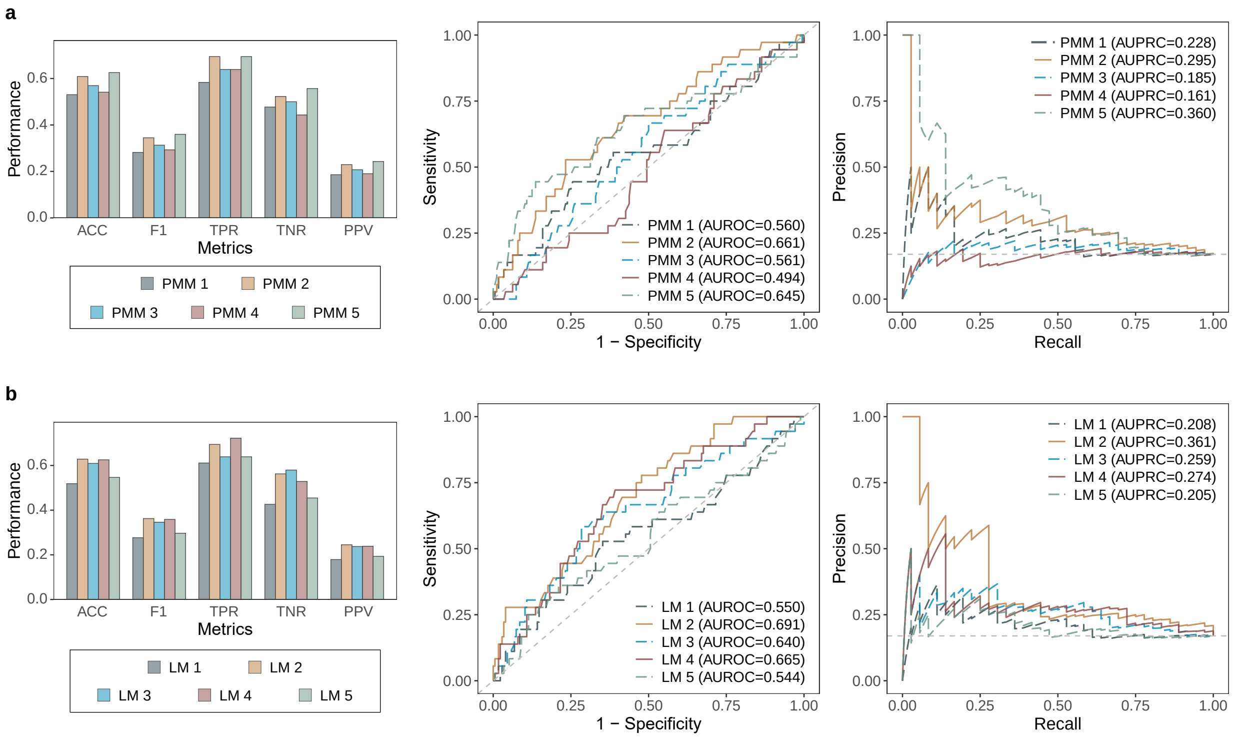

**Fig. S6** The performance of CKD prediction on the KORA multi-omics dataset using each individual imputation of methods PMM (**a**) and LM (**b**). Methods are evaluated on $\mathcal{D}^{\text{test}}_{v=v2,t=t2}$ ($N = 212$, $N_{positive} = 36$, $N_{negative} = 176$), under $obsNum = 0$. Models are trained using the BRF algorithm with identical hyperparameters and evaluated using LOOCV. The barplot (left) shows multi-metric performance. The dashed lines in the ROC (middle) and PR (right) curves represent the performance of a hypothetical model with no predictive capability.
